# Description, measurement, and automatic classification of the *Plasmodium berghei* oocyst morphology during early differentiation

**DOI:** 10.1101/2020.09.15.299024

**Authors:** Benito Recio-Tótoro, Adán Guerrero, Humberto Lanz-Mendoza

**Affiliations:** Centro de Investigaciones Sobre Enfermedades Infecciosas, Instituto Nacional de Salud Pública (INSP), Cuernavaca, Morelos, México; Instituto de Biotecnología, Universidad Nacional Autónoma de México (UNAM), Cuernavaca, Morelos, México

**Author notes:** Corresponding author: Humberto Lanz Mendoza.

**Keywords:** *Plasmodium*, took, oocyst differentiation, classification

## Abstract

After colonization of the mosquito midgut by the malaria parasite, *Plasmodium* differentiates from an invasive, motile ookinete to a multiplicative, sessile oocyst. Despite their importance in establishing the infection and increasing its population, the morphological transformation associated with these changes in function has been scarcely explored. Oocyst differentiation begins with the formation of a spherical protrusion near the center of the crescent-shaped ookinete. As this protuberance grows, it engulfs the content of the two distal ends, thus rounding the cell. In this work, scrutinized observations of the overall changes in shape, coupled with the migration of the malaria pigment granules and the nucleus into the protuberance, revealed that the movement of the cell content happens in an anteroposterior manner. The resulting data, formalized as morphometric measurements, led to the identification of 5 transitional stages and to the development of a computer training algorithm that automatically classifies them. Since cell differentiation has been associated with redox fluctuations, the classification algorithm was tested with parasites stained with a glutathione-specific fluorescent probe, revealing a redox modulation during differentiation.

## Introduction

Malaria is a burden to many tropical and subtropical regions of the world, causing huge economic losses and the death mainly of children(Sachs and Malaney, 2002). In order to be transmitted to a mammalian host, the malaria parasite must venture into a journey through the mosquito tissues. For this reason, strategies for preventing the disease are focused on mosquito control, while novel approaches also focus on avoiding the mosquito to become infected(Blagborough et al., 2013). Despite receiving little attention compared to the vertebrate parasite phases, the study of the mosquito phases have had a long but sparse series of discoveries that date back to when Ronald Ross first observed the malaria parasite developing in the mosquito(Ross, 1897). Since then, a handful of morphological and structural studies regarding the parasite development within the mosquito have accumulated. It is well known the structural differentiation of gametocytes into gametes, the differentiation of the zygotes into ookinetes, as well as the development of the oocyst into sporozoites, but the knowledge about the differentiation of the ookinetes into oocysts is minimal. This comes with no surprise, as the population of ookinetes encounters a serious bottleneck that often reduces the amount of surviving oocysts to a handful(Smith et al., 2014). The development of *in vitro* culture methods that can support oocyst development has greatly enhanced the studies of these elusive sporogonic stages(Al-Olayan et al., 2002), such as the identification of a transitional stage between the ookinete and the oocyst called the transforming ookinete (took)(Carter et al., 2007).

Within 15 minutes of ingestion of a *Plasmodium*-infected blood meal by the mosquito, the gametocytes differentiate into gametes, and fecundation occurs. The resulting zygotes have to escape the digestion of the blood bolus and the maturation of the peritrophic matrix the mosquito secretes to protect against parasites(Smith and Jacobs-Lorena, 2010). To achieve this, the spherical zygotes differentiate into a highly specialized form of motile zygote or ookinete(Angrisano et al., 2012; Janse et al., 1985; Sinden, 2002; Vlachou et al., 2006). During this differentiation, two rounds of meiotic divisions without karyokinesis occurs, while a three-layered membrane called the pellicle starts to grow as a protrusion from the zygote(Sinden et al., 1985; Sinden et al., 1987). This protrusion becomes the anterior pole of the ookinete, which harbors the apical complex, a structure containing the organelles required for locomotion and invasion of the midgut epithelium(Canning and Sinden, 1973). This retort-shaped form of the zygote continues to develop by growing the protrusion, where the malaria pigment, inherited from the macrogametocyte and which is composed of the insoluble hemozoin as a result of hemoglobin digestion(Müller, 2004), moves towards this protuberance. Then the bulbous posterior end gets absorbed, and the nucleus migrates to take a centro-posterior position. The pigment granules left behind follow the nucleus and, in conjunction with the pigment granules that moved before the nucleus, accommodate at both sides of it but are still disperse. In the mature ookinete, the anterior pole enlarges and the pigment granules condense into dense agglomerates(Ghosh et al., 2010; Janse et al., 1985). The mature ookinete is surrounded, from the anterior pole to the posterior, with the pellicle, which constitutes the locomotion machinery(Tremp et al., 2008). Then the ookinete glides out of the blood bolus, crosses the peritrophic matrix, and invades the midgut epithelium(Kan et al., 2014; Vlachou et al., 2004; Zieler and Dvorak, 2000).

Our knowledge of what happens next in regards to the morphological transformation is scarce. It is known that once the ookinete crosses the midgut, reaching the basal lamina that surrounds the organ, an astonishing differentiation begins that transforms the highly specialized invasive ookinete into a trophic sporozoite-producing oocyst, a process that takes up to 15 days to complete. Within the virtual space between the basal labyrinth of the epithelial cells and the basal lamina to which it firmly attaches, the ookinete starts its differentiation into an oocyst by producing a single-layered membrane protrusion in the convex side of the ookinete, which gives the took a snail-like appearance(Carter et al., 2007; Syafruddin et al., 1992; Tremp et al., 2008). The protrusion continues to grow until the spherical young oocyst is formed.

In the young oocyst, the apical complex and the inner membrane complex that comprises the pellicle are reabsorbed(Baton and Ranford-Cartwright, 2005; Canning and Sinden, 1973). Then sporogony begins, a process in which each of the 4 haploid genomes undergoes about 8 rounds endomitosis, giving rise to a population of thousands of genome copies within a single enlarged and convoluted nucleus. While this happens, a bi-layered capsule is formed by the fusion of the oocyst wall with the basal lamina. The mitochondria and apicoplast proliferate, and a series of invaginations of the plasma membrane start to coalesce with small peripheral vacuoles, surrounding the now individual oval nuclei and forming the sporoblast(Baton and Ranford-Cartwright, 2005). The sporoblast grows up to 50-60 µm in diameter while the sporozoites are being assembled and covered with laminin(Aly et al., 2009; Baton and Ranford-Cartwright, 2005; Nacer et al., 2008). Once the sporozoites mature, the oocyst releases them into the haemocel through fenestrations that form in the oocyst capsule(Aly et al., 2009; Baton and Ranford-Cartwright, 2005; Sinden, 1974).

In this work we examined through light microscopy and image analysis the morphological changes that occur during the early oocyst differentiation *in vitro* and compared them with the differentiation of previous and posterior developmental phases of the parasite. This conduced to the finding of common patterns in differentiation and the establishment of 5 readily recognizable steps in the transformation of the tooks. In addition, we offer the community an algorithm in the form of a macro to measure several morphological characteristics of the tooks and to classify them in an unsupervised manner. The macro uses images of the parasite and its nucleus fluorescently labeled, as well as bright field images to segment or isolate the parasite from the image and take morphological measurements. Several hundreds of parasites were measured to build a morphometric database, against which the parasites are compared to, thus assigning them a differentiation stage based on the proximity of the measurements with other parasites.

## Results

### Changes in oocyst morphology during early differentiation follow a specific succession of events which led to the recognition of 5 transitional stages

To study and describe the early development of the oocyst, *in vitro* cultured *P. berghei* ookinetes were purified in extracellular matrix (ECM) gel-coated coverslips and placed in oocyst medium to allow for differentiation. Upon observing thousands of oocysts differentiating, it was recognized that the transformation process follows a similar pattern as in the differentiation of ookinetes(Janse et al., 1985; Sinden et al., 1987) and hepatic trophozoites(Jayabalasingham et al., 2010; Kaiser et al., 2003). Although another classification of ookinete differentiation exists(Ghosh et al., 2010; Siciliano et al., 2020), the classification of Janse, et al. (1985)(Janse et al., 1985) was instead used for comparison since it takes into account the migration of the malaria pigment granules. The morphological characteristics used to establish the proposed classification were the emergence and growth of a protuberance in the ookinete, in conjunction with the displacement of the nucleus and pigment granules towards this ongrowing protuberance (Fig. 1).

**Figure 1.**
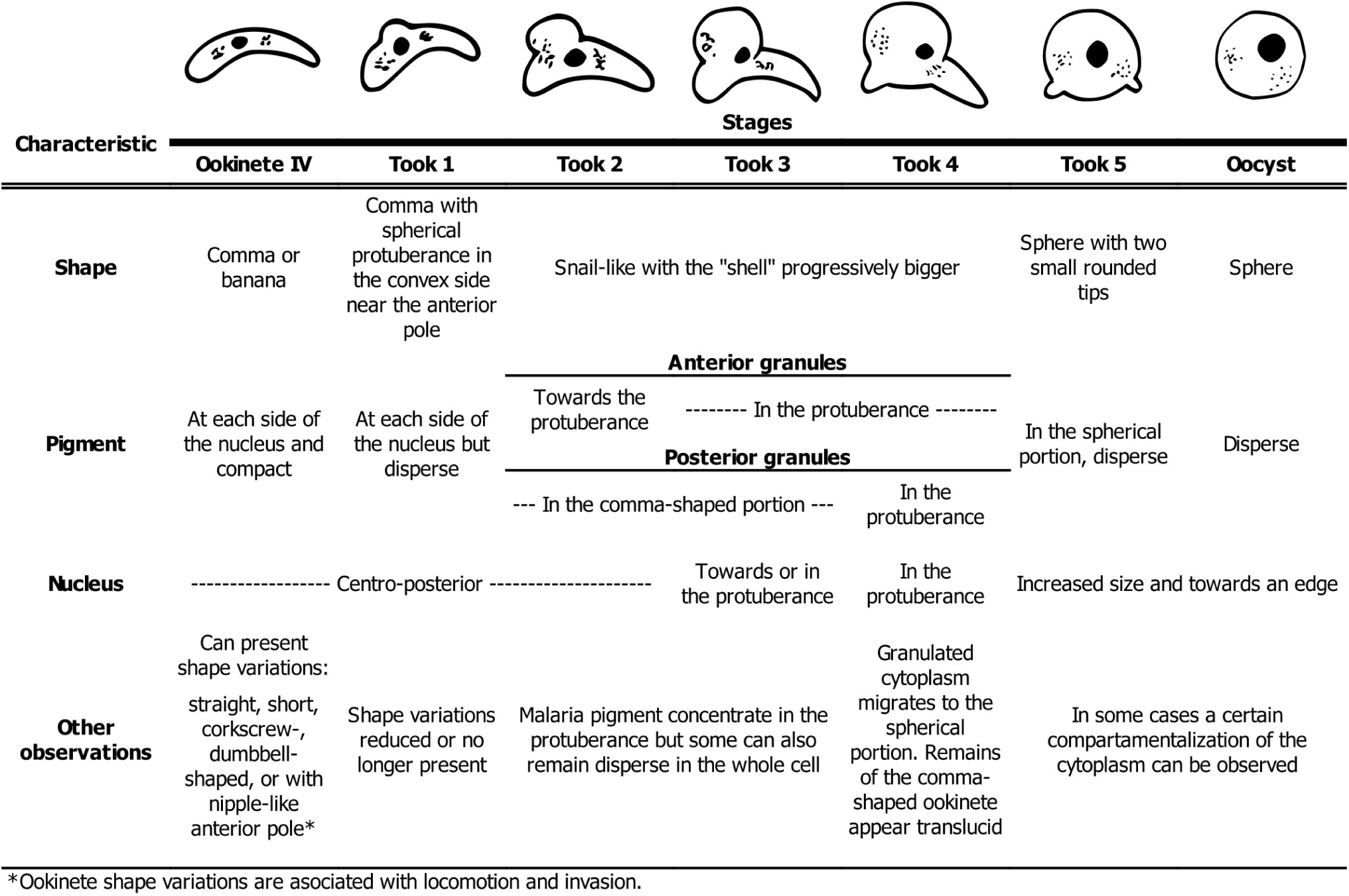
Main characteristics used to establish the classification. The drawings represent the consensus morphology after visual scrutiny of thousands of differentiating parasites visually.

The earliest morphological sign indicating that the ookinete began differentiating is the dispersion of the hemozoin pigment into small granules, which were previously congregated as dense granules adjacent to one or two sides of the nucleus. These ookinetes were still regarded as stage VI ookinetes since, in some parasites, the dispersion of the pigment granules happened at par with the formation of the protuberance. Therefore, the formation of the spherical protuberance near the anterior pole and most commonly in the convex side of the ookinete, is the first reliable morphological sign that differentiation has started (took 1). As this protuberance grows, the anterior pigment granules move to this protuberance (took 2) followed in succession by the nucleus (took 3) and the posterior pigment granules (took 4). As this happens, the ookinete is gradually transforming from a banana-shaped cell into a snail-like cell. Then to a sphere with two small protrusions of what previously were the anterior and posterior ends of the ookinete (took 5), and finally into a sphere (oocyst).

Interestingly, while oocysts are sessile, the tooks are able to glide even at later stages of differentiation (Movie 1), implying that the inner membrane complex and the apical complex are still functional in the remaining portions of what was the ookinete. The spherical part of the tooks, being a single membrane, does not have an inner membrane complex(Tremp et al., 2008) and cannot contribute to moving the parasite.

### Morphometric characterization of the early oocyst differentiation by image analysis

To strengthen the proposed classification, these otherwise subjective observations were formalized as morphometric measurements of the differentiation process. This was achieved by image analysis (Supplementary Fig. S1, see Supplementary Information: Took classification macro), for which fluorescent images of GFP-expressing or stained parasites (see Methods section) were used to obtain the descriptors of shape and size. The nucleus of the parasites was also fluorescently stained to determine its position and size, and bright-field images were used to describe the position, amount, and size of the malaria pigment granules during differentiation (Fig. 2A to 2C). To perform the measurements, the parasites, nuclei, and malaria pigment must be segmented from the images (Fig. 2D). This was carried out in ImageJ taking advantage of the filters and operations this software offers (Supplementary Figs. S2 and S3, see Supplementary Information: Segmentation of parasites, nuclei, and malaria pigment).

**Figure 2.**
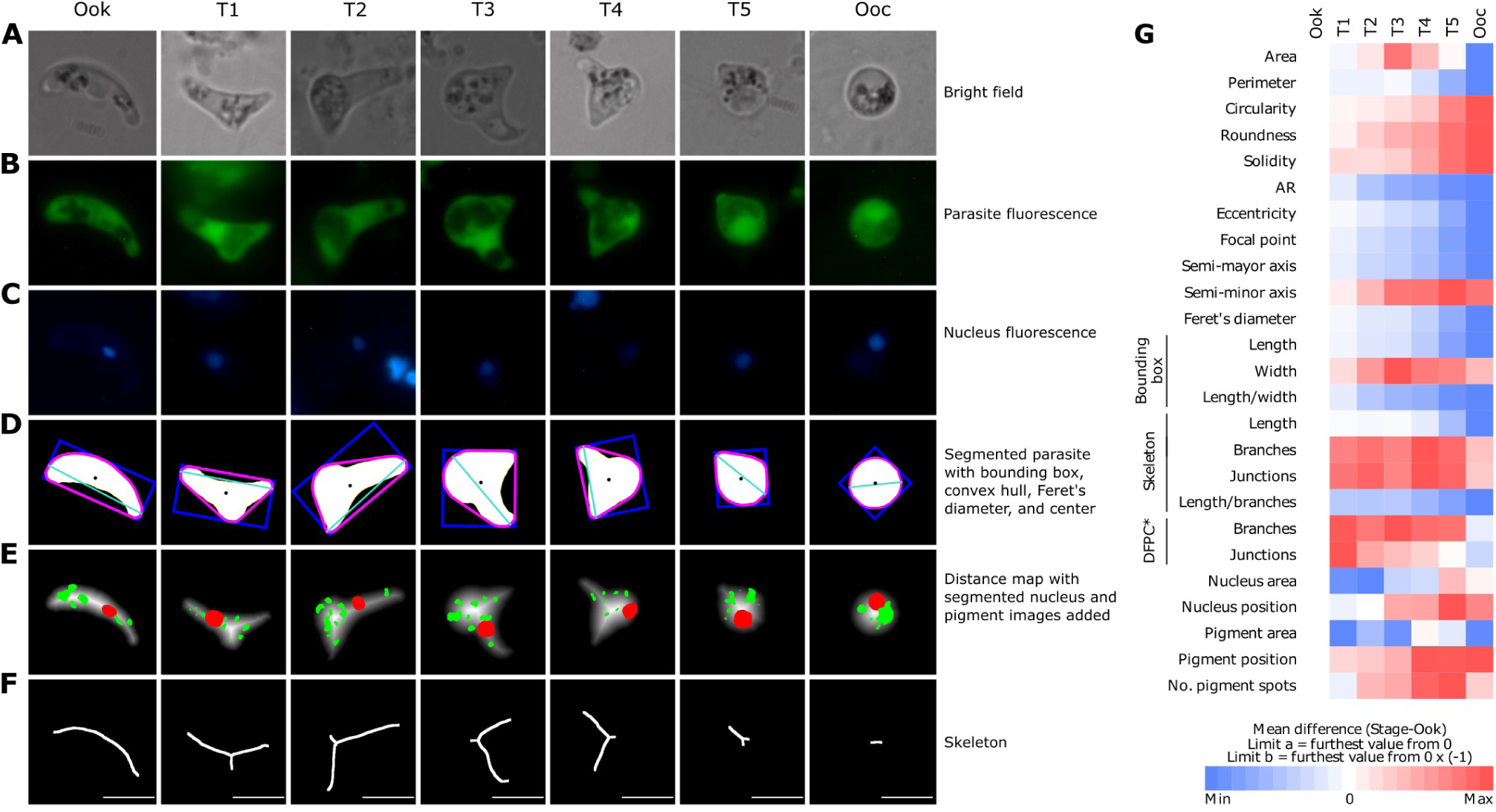
Example images for each differentiation stage and their processing to extract the morphological descriptors of the parasites. **(A)** Bright field, **(B)** parasite’s GFP fluorescence, and **(C)** nucleus’s Hoechst fluorescence images were used to obtain the **(D)** segmented parasite, **(E)** distance map, and **(F)** skeleton images from which the measurements were taken. The segmented parasite (D) image was obtained by applying an edge detection filter on the parasite’s fluorescent (B) image, and was used to obtain the measurements of size, including the bounding box (blue), the adjustment to a circle, to an ellipse, the convex hull (magenta) used to calculate the solidity, the Feret’s diameter (turquoise), and the parasite’s center (black dot). Similarly, the nucleus and pigment granules were segmented from the nucleus fluorescence (C) and bright field (A) images respectively by thresholding algorithms. These images were then used to determine the position of the nucleus (red) and the pigment granules (green) within the parasite by overlapping them on a distance map (E) image of the parasite. In turn, the distance map image was obtained from the segmented parasite’s (D) image, where each foreground pixel is replaced with a gray value proportional to its distance from the nearest background pixel. The skeleton (F) image was also obtained from the segmented parasite’s (D) image by a thinning algorithm that reduces the shape to a single pixel wide line. **(G)** Heat map representation of the changes in the descriptors throughout the stages with respect to the ookinete. Minimum and maximum ranges were calculated independently and normalized for each descriptor. The changes represent the averages of 706 parasites.

A total of 25 descriptors were found to cover broadly the characteristics depicted in Fig. 1. Since the differentiating ookinete transitions from an ellipse-like cell into a circle during differentiation, it was useful to measure how the intermediate stages adjusted to a circle and an ellipse. The latter involved three measurements: the eccentricity, the semi-minor axis, and the semi-major axis. From these other three measurements were obtained: the roundness of the parasite, the aspect ratio (AR), that is, the ratio between the major axis and the minor axis, and the distance from the center of the parasite to the focal point or linear eccentricity. The solidity of the parasites was also determined, which is a measurement of the concave curvatures of the parasites, like the concave portion of the ookinete and the transitions between the protuberance and the main shaft. Another measurement was the longest distance between any two points of the parasites, or Feret’s diameter, which approximates the maximum length.

The changes in size were tracked by measuring the area and perimeter of the differentiation stages, but also by measuring the length and width of the fitted bounding box that encloses the parasites. Additional measures included the analysis of the skeleton of the parasite, that is, a single-pixel-width line representation of the original shape (Fig. 2F). The skeleton image allowed the detection of the protuberance in the tooks by the appearance of a branch. Since this protuberance forms in the anterior portion of the ookinete and slowly displaces to the center as it grows, its position was determined by calculating the distance from the junction of the branches, and from the average distance of the branch tips to the center of the parasite.

The relative position of the nucleus and pigment was obtained through a distance map image of the parasite (Fig. 2E), where the foreground pixels are replaced with grayscale values according to their nearest background pixel. Therefore, the mean gray intensity of the nucleus and pigment selections added onto the distance map reflects their position. Finally, in addition to the area of the nucleus and pigment, the number of pigment spots was also measured (Supplementary Fig. S4, see Supplementary Information: Image processing for measurement of descriptors).

The results of 706 measured parasites are depicted in Fig. 2G. Descriptive and inferential statistics of the descriptors are in the Supplementary Figs. S5 to S7 and in the Supplementary Table S1, respectively. The area of the differentiation stages increased steadily to the took 3, then decreasing and reaching its lowest value in the oocysts (Fig. 2G). The perimeter, AR, eccentricity, focal point, Feret’s diameter, bounding box length and skeleton length decreased during differentiation at different rates, while the circularity, roundness, and solidity increased (Fig. 2G). The bounding box width, being also a reflection of the protuberance size, behaved similarly to the area, increasing to the took 3 and then decreasing. On average, the took 3 represents the largest differentiation stage in terms of area, and it is when the protuberance almost reaches its maximum size, while the two distal ends are still the same size as in the ookinete. From the took 4, the two distal ends start to get absorbed by the spherical portion, which also begins to show in the concave curve of the parasite. Although the oocysts apparently are smaller than the ookinete, its volume is 4.4 times larger (ookinete = 20.94 µm^3^, oocyst = 91.93 µm^3^).

The nucleus and pigment position also increased, meaning that they gradually moved towards the periphery of the cell (Fig. 2G). The number of branches in the skeleton increased from the ookinete to the took 1, remaining constant until the oocyst where it decreased again (Fig. 2G). For the ookinete and the oocysts, being close to an ellipse or a circle, respectively, their number of branches should be one; however, deviations from these shapes, especially irregularities on the edges, can introduce noise that makes them have more than one branch (Fig. 2G). The length of the semi-major and semi-minor axis behaved inversely to one another as expected from the other ellipse-fitting measurements (Fig. 2G). The same is for the number of junctions in the skeleton, which follows a similar pattern to the number of branches.

Confirming the qualitative observations, the quantification of the pigment and nucleus position indicated that the pigment movement, which significantly changed in the took 2 in relation to the ookinete, precedes the nucleus movement, which started to move in the took 3 (Fig. 2G; Supplementary Fig. S7 and Table S1). The number of pigment spots also changed significantly during differentiation, but much later than the observed and described in Fig.1. This is because the segmented pigment granules obtained after image processing were smaller and lower in number compared to the manual segmentation (Supplementary Fig. S3). Thus, only big and sufficiently separated pigment spots were detected individually. Contrary to the observations of Fig. 1, the nucleus area did not change (Fig. 7, Supplementary Table S1), which exemplifies the importance of making quantitative observations.

### Automatic classification of the oocyst differentiation stages through morphometric analysis and a machine learning algorithm

Relying on qualitative observations to classify the differentiation stages can result in wrong stage assignments, especially in instances where a parasite has unclear morphological traits or transit between two differentiation stages. This has happened before in regards to the classification of the ookinete differentiation stages, where the succession order of the transitional stages has been revisited based on more accurate observations (see (Ghosh et al., 2010) and the correction in (Siciliano et al., 2020)).

A way to consistently classify the tooks is to delegate the task to a computer, thereby eliminating the subjectivity of the observer. This was done by analyzing the morphometric data of the parasites through principal component analysis (PCA), which spread the parasite’s descriptors on a multidimensional space to address how much they contribute to explain the variance of the dataset. In this sense, the use of correlated or inversely correlated variables, like the adjustment to a circle or to an ellipse is what spreaded the parasites in the PCA space the most. A starting collection of 28 parasites, 4 for each differentiation stage, with their descriptors computed as in Fig. 2, and previously classified to establish the desired nominal categories, served as seed for the training database. Newly measured parasites, classified by the trainer and by the algorithm, were then added to the training database to extend it. The classification of a parasite into a stage is, on one hand, performed by the trainer following the description presented in Fig. 1; and on the other hand, computed by predicting the PC’s from its morphometric data and comparing them with the centroids of the data cluster of each stage, returning the closest centroid as the predicted stage of the parasite-to-classify. A total of 706 parasites were classified in this manner (Fig. 3A, Supplementary Fig. S8; see Supplementary Information: Automatic classification).

**Figure 3.**
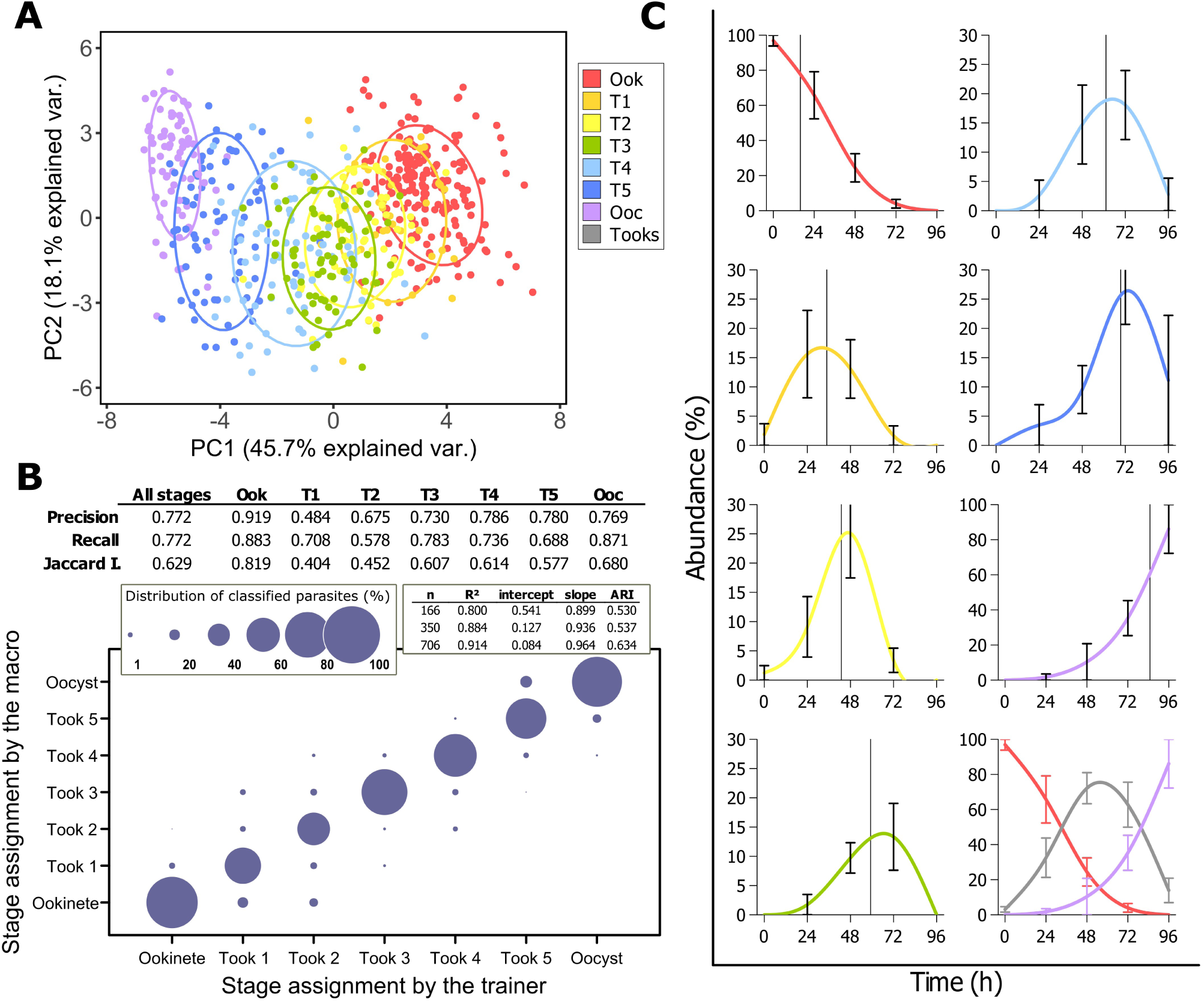
Principal component analysis and automatic classification of the differentiation stages. **(A)** Dispersion of the data of 706 parasites within the first two principal components. The ellipses represent the 68% confidence interval. **(B)** Comparison between the classification of the parasites by the trainer and by the macro. Precision, Recall and Jaccard index shown in the table were calculated for each stage after all parasites were classified. Goodness of fit measurements are resumed in the inset table for different sizes of training databases. Starting from a seed of 28 parasites, each new parasite was classified by the experimenter and by the algorithm simultaneously and added to the training database to extend it. **(C)** Once the whole set of parasites were classified, the percentage of each stage per day was obtained and plotted as transformation kinetic curves. Ook = ookinete, T1 = took 1, T2 = took 2, T3 = took 3, T4 = took 4, T5 = took 5, Ooc = oocyst, Tooks = T1 to T5 grouped together. The vertical line in each curve represents the mean of the abundance distribution. Error bars represent the standard deviation of the mean.

The prediction results in comparison to the visual assignment made by the experimenter are shown in Fig. 3B. To evaluate the accuracy of the predictions, the precision, recall, and Jaccard Index were determined, as explained in the methods section. In all three coefficients, higher values indicate higher performance by the algorithm, where a value of 1 indicates perfect performance. The precision tells the proportion of correct assignments of a stage among all instances where that stage was assigned, regardless if it was incorrect. The recall tells the proportion of correct assignments of a stage among all instances where that stage should have been assigned. Since wrong assignments for one stage imply a wrong assignment for another stage, the precision and recall are equal when all stages are considered. The Jaccard index is a measure of similarity that tells how similar the assignments made by the trainer and by the algorithm are.

Additional measures of goodness of fit were the R^2^, the slope, and the intercept of the observed vs. predicted curve, represented as a bubble chart (Fig. 3B). For a good fit, the R^2^ and the slope would approximate to 1 and the intercept to 0. The Adjusted Rand Index (ARI), ranging from −1 to 1, where 0 is random, and 1 is total similarity, was also calculated. This index also tells the agreement of two datasets, but, in contrast to the Jaccard Index, it considers the chance of random correct assignments. The performance of the algorithm improved as the database increased as shown in the inset table of the bubble chart and in the Supplementary Fig. S9.

In general, the agreement between the trainer and the algorithm was above 0.6, indicating that in most instances the parasites were classified by the algorithm as expected. Exceptions were found at the took 1, the took 2, and to a lower extent at the took 5. The main difference between the ookinete and the took 1 in terms of the measured descriptors, is the appearance of a junction with its branch in the skeleton. Sometimes the protuberance in the took 1 is small enough, or the parasite is positioned in such a way that this branch does not appear in the skeleton, thus, the parasite is classified as ookinete instead of took 1. Something similar occur with the took 1 and 2, and between the took 2 and 3. All these stages presented the fewer number of significant changes in their descriptors (Supplementary Table 1). Most of the differences found between the took 1 and 2, and between the took 2 and 3 reside in their ellipse fitting measurements and in the width of the fitted bounding box, which represents the growth of the protuberance. Interestingly, these stages are the most difficult to classify visually, so it may be possible that the trainer is the one mistaken when classifying them.

The results of the classification allowed the formation of differentiation kinetic curves for each one of the stages. As seen in Fig. 3C, the differentiation curves of the tooks are arranged accordingly to its differentiation stage, (e.g. the mean peak abundance of tooks 1 is previous to tooks 2, then tooks 3, tooks 4 and finally tooks 5), which is another indication of the correct classification of the stages.

### Fluorescence analysis of mBCl-stained parasites reveals changes in GSH levels during differentiation

As a proof of concept for the classification, the macro was tested with parasites stained with the glutathione (GSH) specific probe monochlorobimane (mBCl). This probe is coupled to GSH by means of the GSH S-transferase enzyme forming the GS-bimane adduct and increasing 10-fold its fluorescence. GSH is a γ-tripeptide containing cysteine involved in the redox homeostasis of the cell and, because of its high concentration and ubiquitous presence in the cytoplasm and organelles of most eukaryotes, it is well-suited for staining cells relatively uniformly(Keelan et al., 2001).

It is known that differentiating cells enter a state of redox imbalance where the total GSH content is decreased(Toledo et al., 1995). Transcriptome analysis has shown oscillations in the expression of the enzymes that synthesize and recycle GSH in the erythrocytic development of the parasite(Bozdech and Ginsburg, 2004), suggesting a decrease in GSH content in the phases that are differentiating. The decrease in GSH content is involved in the establishment of more oxidizing conditions in the cell that allows for reactive oxygen species (ROS) accumulation and redox signaling(Hernández-García et al., 2010; Schafer and Buettner, 2001). Furthermore, GSH itself is involved in signaling and protein protection through glutathionylation, effectively decreasing the free GSH in the cell (Berndt et al., 2014; Cooper et al., 2011). In addition to the GSH measurement, an attempt to measure reactive oxygen species (ROS) through the differentiation with the ROS-sensitive fluorophores H_2_DCFDA and CellROX was unsuccessful, as no signal could be detected in any of the stages.

First, the mBCl fluorescence was characterized in ookinete cultures by kinetic curves with different concentrations of the probe, showing that the fluorescence presented a wide linear detection range (Fig. 4A). For further experiments, the parasites were stained for 30 min with an mBCl concentration of 160 µM to allow a wide working margin. Then, the fluorescence intensity of the GS-B adduct was measured in the stages using the algorithm developed herein. It was found that the GSH content decreased slightly in all stages through time of culture (slope of the curves: Ook = −253 fluorescent units per day (FU/day), T1 = −224 FU/day, T2 = −136 FU/day, T3 = −474 FU/day, T4 = −285 FU/day, T5 = −298 FU/day, and Ooc = −136 FU/day). This decrease, however, was not significant (ANOVA comparing days per stage with the lowest *p* value of 0.21 found in the tooks 5), but enough to introduce noise when the GHS content between days was averaged. Therefore, the GSH content was normalized between culture times using the slope of the curve as the normalizing factor. The fluorescent signal was found to concentrate in the nucleus, its surroundings, and in the apical complex, indicating a higher GSH content in these structures. Also, the regions containing the malaria pigment appear darker, meaning that the pigment eclipsed the signal or that the probe was excluded from these zones (Fig. 4B).

**Figure 4.**
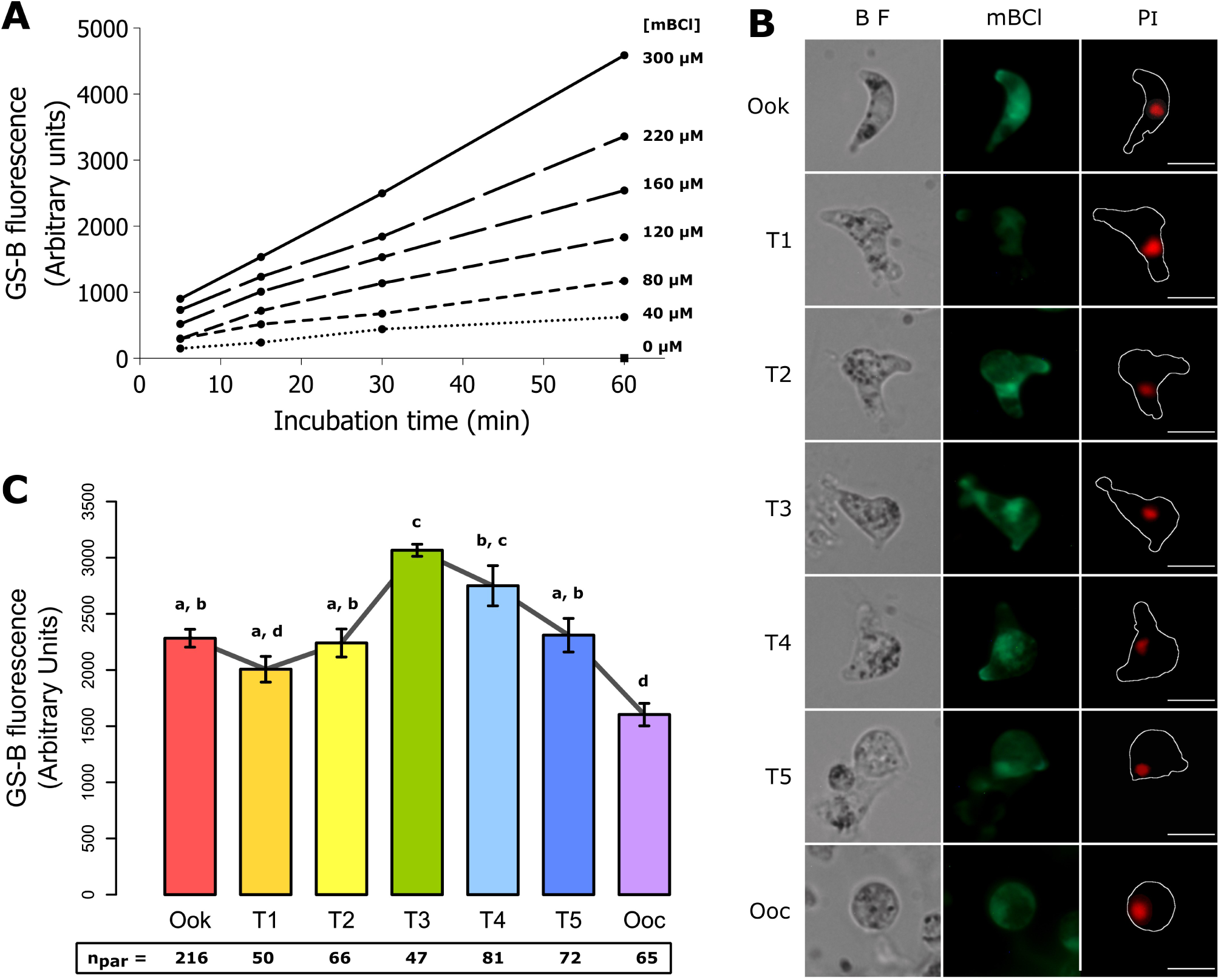
Measurement of the fluorescence intensity of the GS-B adduct during differentiation. **(A)** mBCl concentration curve and incubation time in ookinetes. Results of 2 independent experiments. **(B)** GSH content in the differentiation stages. Data represent the mean and standard error of the mean of 3 independent experiments analyzed by ANOVA and Tukey *post hoc* test, bars with different letters are statistically significant at an α level of 0.05. n_par_ = number of parasites sampled. **(C)** Representative images of mBCl and PI-stained parasites. Ook = ookinete, T1 = took 1, T2 = took 2, T3 = took 3, T4 = took 4, T5 = took 5, Ooc = oocyst, BF = bright field, mBCl = monochlorobimane, PI = propidium iodide.

Interestingly, the average GSH content varies significantly during differentiation (Fig. 4C). First, there is a slight non-significant decrease of GSH in the took 1, increasing afterward until it reaches its highest content in the took 3 with 1.3 times more GSH in comparison with the ookinete. The GSH content then starts to decrease again, reaching the lowest content in the oocyst with 1.9 times less in comparison with the took 3. These data suggest that there might be indeed redox-associated signaling during oocyst differentiation, in agreement to what it has been observed in the differentiation of other phases of the malaria parasite.

## Discussion

More than 30 years have passed since the morphology of the differentiating ookinete was described(Janse et al., 1985; Sinden et al., 1985; Sinden et al., 1987). Around this time, work was also done regarding the oocyst development(Canning and Sinden, 1973; Syafruddin et al., 1992), and the differentiation of the sporozoite into hepatic trophozoite(Meis et al., 1983; Meis et al., 1985). These seminal works paved the way for more recent efforts that are still finding new morphological features of the parasite’s sporogonic development(Carter et al., 2007; Jayabalasingham et al., 2010; Kaiser et al., 2003). Despite this, the detailed progression of the early oocyst differentiation had not been determined before.

In this work, the morphological changes in the early oocyst development were characterized by light microscopy of *in vitro* cultured parasites. In addition to the qualitative description, the differentiation process was also followed through image acquisition and analysis. This allowed the obtainment of quantitative measurements that had hitherto not been performed and, ultimately, to the development of an ImageJ macro to automatically sort the differentiation stages. The differentiation process is a gradual and continuous progression of shape changes and organelle movements as shown in Figs. 1 and 2, thus, no real stages actually exist. However, establishing limits and organizing the observations into discrete classes’ aids in structuring the understanding of how differentiation happens and eliminates the subjectivity one can encounter during experimentation, leading to consistency and reproducibility. In the algorithm presented here, this gradual progression in changes was categorized into 7 differentiation stages including the ookinete and the oocyst, where a parasite is classified in relation with its proximity to the centroids of the data clusters of previously classified parasites. In this way, visually hard to classify parasites due to mixed characteristics with previous and posterior stages, are assigned to a stage consistently and objectively.

Despite the measurements behaved as a continuum, the rate of change was not always constant at specific points during differentiation. The fact that some descriptors changed at slower rates while others at higher rates, or even changed only at particular points during differentiation, is what allowed classification. The use of several descriptors is also critical as it guarantees that at least one descriptor is useful for distinguishing two or more stages, as happens between the took 1 and 2, and between the took 3 and 4, in which different descriptors are the ones useful.

The differentiation rates obtained here are comparable to those obtained in other studies(Al-Olayan et al., 2002; Azevedo et al., 2017; Carter et al., 2003; Carter et al., 2007; Porter-Kelley et al., 2006; Syafruddin et al., 1992), with the addition that we now know the kinetic differentiation curve for each differentiation stage plus the means to identify them consistently. This could, for example, lead to the identification of critical steps during differentiation, as recently found for the ookinete differentiation(Siciliano et al., 2020).

Carter and co-workers (2007)(Carter et al., 2007), pointed out the differences between the differentiation of the ookinete and the differentiation of the oocyst. In this work, we rather focused on the similarities to establish a classification, which also resembles the differentiation of sporozoites into hepatic trophozoites considerably(Jayabalasingham et al., 2010; Kaiser et al., 2003; Meis et al., 1985). However, important differences still reside, which may help to unveil molecular and metabolic processes.

The differentiation of the oocyst can be seen as the same as the differentiation of the ookinete but in reverse. While the membrane of the zygote is single-layered, the protrusion that starts to grow in the retort is three-layered(Sinden et al., 1987). Then, as the protuberance grows, a portion of the malaria pigment starts to move towards this protuberance, followed by the nucleus, which, in the zygote, is located towards the periphery of the cell; finally the other portion of the malaria pigment follows the nucleus. Both the portion of pigment that moved first, and the other portion that moved last; condense at both sides of the nucleus, which takes a central-posterior position(Janse et al., 1985). Then, as explained earlier, once the ookinete starts to differentiate into an oocyst, the pigment granules disperse again and a single-layered membrane protrusion starts to grow(Carter et al., 2007; Tremp et al., 2008), albeit at the centro-anterior portion and predominantly in the convex side of the ookinete. The anterior pigment granules move to the protuberance as it grows, followed by the nucleus and then by the posterior pigment granules. In the young oocysts, the nucleus again takes a position closer to the periphery rather than centered, with the pigment granules remaining disperse throughout the cytoplasm.

In the hepatic trophozoite differentiation, a single-layered membrane starts to grow in the sporozoite, resembling the oocyst differentiation; however, in a more central location and where the nucleus is. This causes the nucleus to be right in the protuberance as it appears, which can be at either the convex or concave edge of the sporozoite(Jayabalasingham et al., 2010). Since the sporozoite lacks malaria pigment granules, there is no analogy to the differentiation of the oocysts in this regard. Furthermore, similar to the oocyst differentiation, in the differentiating hepatic trophozoite there is a clearance of the specialized organelles the sporozoite uses for locomotion and invasion. However, it was shown by Jayabalasingham and co-workers (2010)(Jayabalasingham et al., 2010) that the inner membrane complex and the micronemes are being discharged into the parasitophorous vacuole. This has not been observed in the differentiating oocyst and indicates differences in organelle recycling, probably due to the parasite location within the host. Since the oocyst differentiation occurs extracellularly, in contrast to the differentiation of the hepatic trophozoite, which is intracellular, reabsorbing the invasion-specific organelles instead of discharging them may prevent recognition by the host. Canning and Sinden (1973)(Canning and Sinden, 1973) reported that the apical complex in the ookinete degenerates and reabsorbs immediately after transformation into oocyst starts. More recently, however, it was found that the apical complex is retained until late in the differentiation process(Carter et al., 2007). This was also indirectly observed in this work (since GSH concentrates in the apical complex and the nucleus), showing that the apical complex still remains in its position in the took 5, disappearing in the oocyst.

In sum, our observations enriched what we knew about the differentiation of the oocyst and furthermore, not only establish common patterns in the differentiation of the sporogonic stages but also point out significant differences; both of which may lead to the development of new hypothesis about the molecular machinery that is in charge of these processes. Though unexpected, the ability of the tooks to continue gliding while differentiating explains why they conserve the apical complex until the oocyst is formed. This machinery may allow the ookinetes to escape the entry zone from where they reached the basal lamina and evade the localized immune response triggered by the apoptotic epithelial cell(Han and Barillas-Mury, 2002; Whitten et al., 2006) while differentiating at the same time.

The changes in GSH content between the stages suggest modifications in the redox status during differentiation. Modifications in the redox status have been regarded as a common hallmark of cell differentiation and other cell destiny determination processes like proliferation and programmed cell death(Aguirre et al., 2005; Schafer and Buettner, 2001). In the parasite, redox changes have been observed in the erythrocytic development(Bozdech and Ginsburg, 2004), in the commitment to sexual development(Beri et al., 2017; Chaubey et al., 2014), and in the ookinete differentiation(Duran-Bedolla et al., 2016). Further studies should be made to address the redox changes during oocyst differentiation, especially since one way in which the mosquito responds to the parasite invasion is by producing reactive oxygen species(Han et al., 2000; Herrera-Ortíz et al., 2004; Lanz-Mendoza et al., 2002; Luckhart et al., 1998). Even more, it is known that the GSH synthesis and recycling is indispensable for the sporogonic phases(Pastrana-Mena et al., 2010; Vega-Rodríguez et al., 2009). The redox changes could be studied, for example, by measuring the NADPH/NADP or GSH/GSSG redox couples, as well as the proteins involved in these processes during differentiation, and identify which stages are more susceptible to an oxidative insult. Knowing the oxidative stress response of the parasite could lead to the finding of new targets for the development of transmission-blocking interventions.

In summary, we present a detailed morphological examination of the oocyst differentiation through light microscopy, categorizing the transformation into five differentiation stages (see Movies 2 and 3 for a demonstration of the macro). Additionally, we provide the means to measure and classify the proposed stages objectively and automatically. Albeit the algorithm was written for processing, measuring, and classifying the early oocyst differentiation stages, it can be easily adapted to perform the same tasks, including the classification, with other developmental phases of the malaria parasite, or even with other cells or parasite species, given a training database is built. It can also serve as a framework to write other algorithms to perform more complex and specific measurements, for example, if the apicoplast, the mitochondria, or any other organelle needs to be tracked during differentiation.

## Materials and methods

### Ookinete purification and oocyst culture

*Plasmodium berghei* ANKA strain clone 2.34 and the derived transgenic line that constitutively expresses the green fluorescent protein (GFP)(Franke-Fayard et al., 2004) were maintained in BALB/c mice for *in vitro* ookinete production as described previously(Rodríguez et al., 2002). Ookinetes were harvested 18 h post culture and purified by affinity using extracellular matrix (ECM) gel as the substrate (in preparation). Briefly, the cultures were incubated on coverslips coated with ECM gel for 4 h to allow ookinete attachment. The coverslips were then washed twice with 1 ml of PBS to eliminate the blood cells and other parasite stages. After purification the parasites were maintained in 1 ml of oocyst medium consisting of Schneider’s *Drosophila* medium at pH 6.8 supplemented with 15.87 mM sodium bicarbonate, 20 mM HEPES, 3.68 mM hypoxanthine, 44 µM *para-*aminobenzoic acid (PABA), 0.2% lipid/cholesterol solution (Gibco), 100 U/ml penicillin, 100 µg/ml streptomycin, 200 µg/ml gentamicin, and 15% heat-inactivated fetal bovine serum(Al-Olayan et al., 2002; Carter et al., 2007). The oocyst medium was renewed after 3 days replacing half the volume with fresh medium.

### Staining

For light microscopy, the parasites were fixed with methanol and stained with 30% Giemsa in water for 10 min. For fluorescence microscopy, the GSH-specific fluorescent probe monochlorobimane (mBCl)(Chatterjee et al., 1999; Keelan et al., 2001) was used. The parasites were stained by placing the coverslips in wells with 1ml of 160 µM mBCl in oocyst medium for 30 min, washed twice with PBS, and fixed in 4% paraformaldehyde and 0.2% glutaraldehyde for 15 min. For staining nuclei, the parasites were washed twice with PBS and incubated at 28°C for 45 min in 1 % Triton X-100 and 20 µg/ml of RNase A. After two more washings, the coverslips were mounted on glass slides with 2 µl of 250 nM propidium iodide in PBS and sealed with nail polish. The nuclei of GFP-expressing parasites were stained with 2 µl of 200 nM Hoechst without fixing. Since ookinetes and early tooks are able to move by gliding motility, it was necessary to mount the GFP-expressing parasites on a drop of 0.2% agarose to immobilize them.

### Microscopy, image acquisition, and fluorescence measurement of mBCl-stained parasites during differentiation

The parasites were observed and photographed in an epifluorescence Leica DM1000 microscope at 100X magnification (NA = 1.25, oil immersion) coupled with a cooled Tucsen TCC-3.3ICE-N color camera and the TSView software version 6.2.4.2. The system was spatially calibrated with the use of micrometer rulers. The fluorophores were excited with a mercury lamp using the following filters: GFP was excited with a 420-490 nm filter and visualized through a 510 nm dichroic mirror (DM) and a 515 nm long pass (LP) filter; mBCl and Hoechst 33258 were excited with a 355-425 nm filter and visualized through a 455 nm DM and a 470 nm LP suppression filter; for PI a 515-560 nm excitation filter was used with a 580 nm DM and a 590 nm LP filter for visualization.

Each day, a coverslip was taken as a sample of the oocyst culture, stained, and 10 to 15 random fields of view imaged. Grayscale bright field images were taken for each filter and 15 s color videos were taken for the fluorophores. For this, the fluorescence shutter was opened 4 s after the recording started and 4 s before the recording ended. The brightest slice of the videos was then extracted for further analysis. The videos were recorded uncompressed in AVI format at 1 frame per second with the following exposure times: GFP 0.50 s, mBCl 0.52 s, PI 0.60 s, and Hoechst 0.40 s. Each parasite registered, therefore, has 4 files: two grayscale images and two videos for each set of filters used.

The total corrected fluorescence was obtained as described elsewhere(McCloy et al., 2014) with modifications. The integrated density of each color component was measured separately and then added. The total integrated density was then subtracted by the product of the area of the parasite, times the average of the mean pixel intensity of three random background measurements.

### Statistical analysis and accuracy of the algorithm

All statistical procedures, tests and graphing were performed in R version 3.6.1 (2019-07-05). Data was tested for normality using the Shapiro-Francia test; when this criterion was met, Student’s *t*-test or ANOVA, followed by Tukey HDS *post-hoc* test was performed. In the case of non-normality, data was tested with the Wilcoxon-Mann-Whitney test or Kruskal-Wallis followed by Dunn’s test.

To determine the performance of the classification algorithm, the agreement between the stage assignment by the trainer and the stage assignment by the algorithm was quantified by determining the true positives (TP), the false negatives (FN), and the false positives (FP) as follows. Each stage can be either classified as the expected stage, having a value of 1, or classified as any other stage except the expected one, having a value of 0. Thus, the possible outcome combinations are:

*TP = S*_*11*_, where the trainer and the algorithm agree in the classification of a parasite.

*FN = S*_*10*_, where the trainer assigns the parasite as stage *x* but the algorithm assigns it as stage *y*. And

*FP = S*_*01*_, where the trainer assigns the parasite as *y* but the algorithm assigns it as *x*.

The rest of the classifications that do not fit in these conditions are the true negatives (*TN = S*_*00*_) so that *∑ TP + ∑ FN + ∑ FP + ∑ TN = n*, or the total amount of classified parasites.

Once having the predictions classified into these conditions, the precision, recall and Jaccard index were calculated as follows(Garcés et al., 2016):

The precision, also referred to as the positive predictive value is defined as:

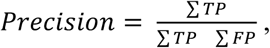

The recall, or sensitivity is defined as:

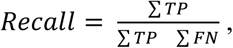

The Jaccard index measures, in the context of this work, the agreement between assignments made by the trainer to those made by the algorithm, and it is defined as:

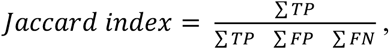

where *∑ TP, ∑ FP*, and *∑ FN* indicate the total number of true positives, false positives, and false negatives respectively.

The Adjusted Rand Index (ARI) measures the similarity of two data clusters considering the chance of random correct assignment. In this case all the possible outcomes between the stage assignments by the trainer vs. the stage assignments by the algorithm. The ARI is defined as follows(Hubert and Arabie, 1985):

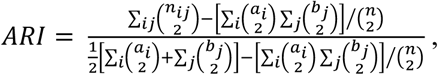

where *n*_*ij*_ are the number of parasites in each combination between the stage assignments arranged in a contingency table (Fig. 3B represents such contingency table), and where *α*_*i*_ is the sum of the rows (the stage assignment by the algorithm) and *b*_*j*_ is the sum of the columns (the stage assignment by the trainer).

## Acknowledgments

The authors want to thank María del Carmen Rodríguez Gutierrez for gifting the two strains of parasites used in this study. The authors also appreciate the useful critical reviews made by Fabiola Claudio Piedras, Mario Henry Rodríguez López, Enrique Salas Vidal, and mostly the reviews made by the students and personnel of the Laboratorio Nacional de Microscopía Avanzada (IBt-UNAM).

## Competing interests

No competing interests declared

## Funding

This work was supported by the Consejo Nacional de Ciencia y Tecnología [487032/277845 to B.R.T].

## Data availability

## Supplementary Information

Movie 1. Differentiating oocysts of *Plasmodium berghei* are still able to glide during transformation. The comma-shaped cells are ookinetes, the snail-like cells are transforming ookinetes (or tooks for short), the circular cells are young oocysts. The parasites were purified from *in vitro* cultures by allowing them to attach to a thin coating of extracellular matrix gel. After washing the contaminant blood cells and other parasite stages, the ookinetes were placed in differentiation medium. Time-lapse images recorded at 1 frame per 2.5 seconds of a 48 h old culture.

Movie 2. Instructions for the installation of the Took Classification Macro. The required programs and plugins can be found in the following links.

ImageJ: /urlhttps://imagej.nih.gov/ij/index.html

R: /urlhttps://www.r-project.org/

RStudio: https://rstudio.com/products/rstudio/

Template Matching and Slice Alignment:

https://sites.google.com/site/qingzongtseng/template-matching-ij-plugin

Morphological Operators for ImageJ:

https://blog.bham.ac.uk/intellimic/g-landini-software/

Took Classification Macro:

https://github.com/BenitoRT/TookClassificationMacro.

Movie 3. Demonstration and usage instructions for the Took Classification Macro. See the documentation of the macro (Readme.txt) for further details.

### Took classification macro

A macro was built in ImageJ *v* 1.53*c* to process and analyze the images of differentiating oocysts stained with fluorescent probes semi automatically. In addition to allow for large batches of images to be processed in a short amount of time compared to a manual processing, the macro also makes possible the obtaining of accurate morphometric measurements of the differentiating parasites and their consistent classification into the 7 proposed stages. The macro is easy to implement, requiring only ImageJ and R statistical software to run. All the procedures and operations used (see following sections) to align, crop, segment, and measure the descriptors and fluorescence intensity of the parasites were programmed in ImageJ’s macro language. The operations performed to classify the parasites were carried out in R. These two programs were linked together via Rserve [1], a socket server application that allows the use of R capabilities in other software, so all the R functions are called within ImageJ. The macro requires that bright field and fluorescence images are taken with each filter used, one for the cytoplasm fluorescence and one for the nucleus fluorescence. In this work, when measuring and comparing the fluorescence intensity between the stages, we found it more reliable to take video instead of pictures of the fluorescence, so the signal is measured at its brightest, nevertheless the macro, is able to process both images and videos. The images/videos have to be named consecutively in ascending order, using leading zeroes or time-stamps.

The macro can be viewed as a three part process (Supplementary Fig. S1). First, if required, the brightest slice of the videos is extracted to obtain the fluorescent images of the parasite and its nucleus. Then the images taken using the different filters are aligned and cropped to leave only one parasite per image. Finally, the parasite, its nucleus and pigment granules are segmented in order to be detected. In a seconds step of the macro, the images are processed to measure the descriptors that are compared against a database for classification in R. For this, ImageJ passes the measurements to R to perform the classification. ImageJ then receives the prediction form R and the parasite is sorted in its corresponding file. During training of the database, ImageJ displays a montage of the parasite and its descriptors, as well as the predicted stage, so the trainer could decide if the prediction was correct or not. The trainer judges the images and assigns a stage to the parasite, which overrides the prediction of the macro. This functionality is by default off but can be turned on if desired. Then, R is called again to save the measurements of the parasites and its differentiation stage alongside the other previous measurements, extending the database. Once all parasites have been classified, the third step of the macro involves going through all images again to measure the fluorescence intensity of the parasite and its nucleus, saving the results in a spreadsheet file. Big batches of parasites do not have to be analyzed in one run. The macro has checkpoints that allow stopping the algorithm and resuming it afterwards from where it was left off. These checkpoints are present after cropping, segmenting, classifying the parasites, and measuring the fluorescence intensity.

The user’s input is required in the three steps of the macro. First, in cropping the parasite for which the images are displayed. After cropping a parasite, the user is asked if more parasites have to be cropped from the same image or to proceed to the next image. Second the user is asked to validate the segmentation of the parasite, the nucleus and the pigment granules. Here, the user can discard the parasite from the analysis if it is not correctly segmented, or segment the parasite by hand if desired. And third, a last step involving the user’s input is for selecting three background zones that will be needed for the fluorescence intensity measurements. Even with the user interventions, the time required to analyze the images is considerably shortened in contrast to performing the analysis solely by hand. This allows for the classification and fluorescence analysis of hundreds of parasites in just hours instead of days. Analyzing a parasite, from the preparation of the images to the measurement of the fluorescence intensity, takes approximately 1 min 25 s. Most of this time is spent in the parts where the user’s input is required. The classification of the parasite takes around 140 ms of computing time with a database of 706 parasites. While the processing, measurement, and generation of the derived images of the parasites takes around 30 s.

This macro is offered as a free download to the research community. For installation, the .zip file has to be extracted into the ImageJ macros folder (Movie 2). For running the macro ImageJ has to be opened and the macro executed form the Plugins -*>* Macros tab (Movie 3). A custom button is also provided for short access.

**Supplementary Figure 1:**
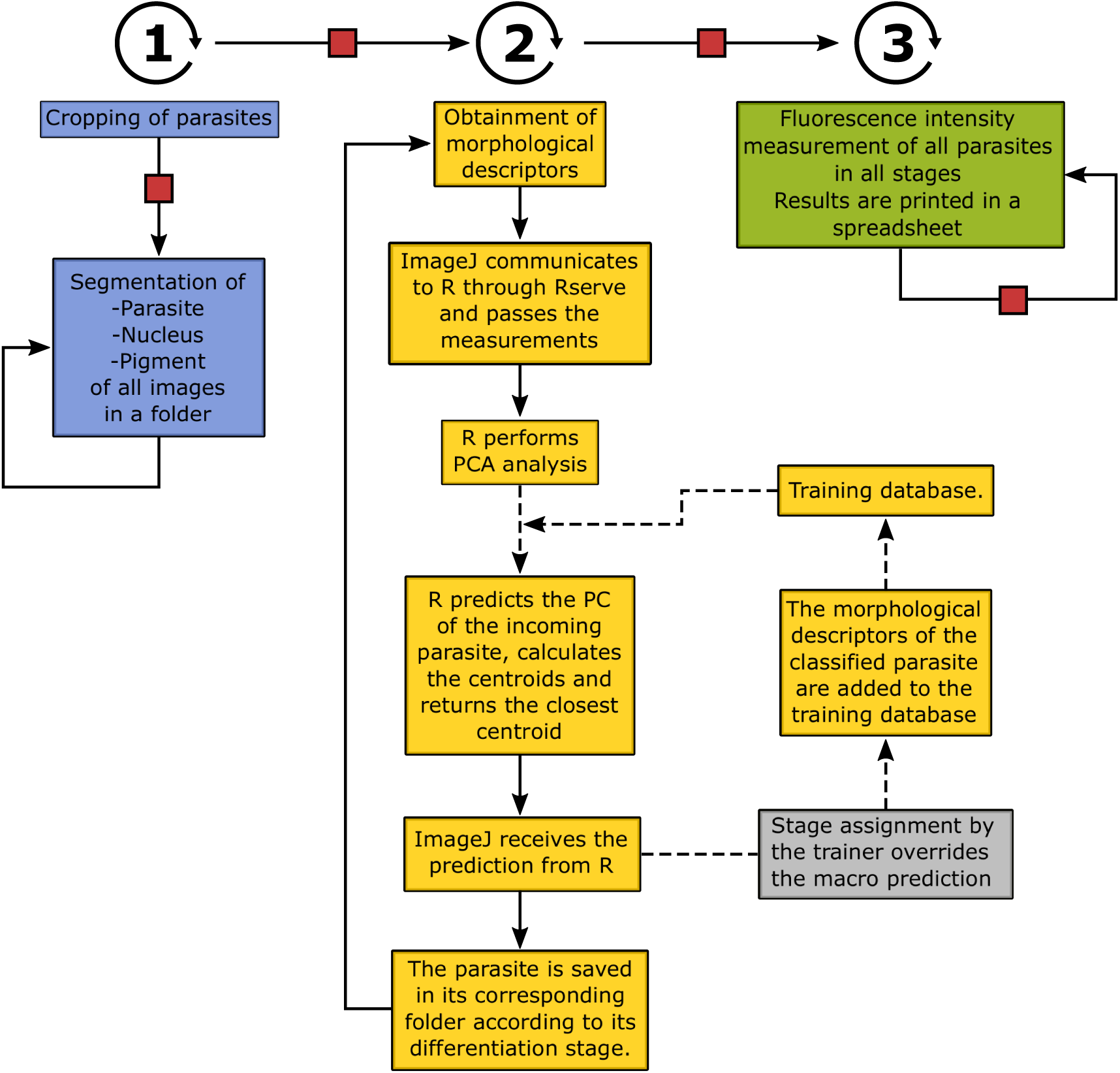
Took classification macro. The macro can be viewed as a three step process (blue, yellow, and green boxes). First the images are cropped and processed for segmentation. In a second step, the measurements of the morphological descriptors are obtained and the parasite classified. If not in training mode, the gray box is bypassed. In a third step, the fluorescence intensity of each parasite at each stage is measured and saved in a spreadsheet. Arrows indicate that the operations are carried out in ImageJ, dotted arrows indicate that the operations are carried out in R. The red squares represent the checkpoints where the macro can be stopped and resumed.

### Preparation of images/videos

The macro needs three images per parasite, one image of the bright field of the parasite, one image of the fluorescence of the parasite, and one image of the fluorescence of the nucleus. The parasite’s and nucleu’s fluorescence images can also be videos. The macro also accepts four images/videos per parasite, where the fourth image/video is the bright field of the filter used to image/record the nucleus. This is in some cases necessary due to the color channel offset or pixel shift that could happen when using different filters, which leads to misinterpretations in the positioning of the nucleus.

After opening the first image of a list of parasites, the macro extracts the brightest slice of the videos for each parasite if needed. Then, and only if needed, the images of each parasite are grouped in a stack and aligned using the ImageJ plugin “Template matching-align slices in a stack”. The stacks are then cropped with the assistance of the user to leave only one parasite per image, simplifying further analysis.

### Segmentation of Parasites, nuclei, and malaria pigment

The steps for performing the segmentation of the parasites, the nuclei, and the pigment granules are explained below and resumed in the Supplementay Fig. S2. For segmenting the parasites, the parasite’s fluorescence image was convoluted with a Laplacian of Gaussian (Mexican Hat) filter with a kennel radius of 20. This detected the edges of the parasites, which were then smoothed, while preserving the edges, with a median filter of radius = 4. The resulting image was then transformed to 8-bit and thresholded to 85, 255. The region of interest (ROI) was then detected using the “Analyze Particles” algorithm with the following conditions: 12-60 µ*m*^2^, circularity between 0.32 - 1.00, including holes, and excluding edges.

For segmenting the nuclei and the pigment, it was quite useful to overlap the parasite segmentation output ROI into the nucleus and pigment images to clear the surroundings. This reduced the complexity of the images and increased the segmentation success for these structures. Then a Gaussian filter of sigma = 3 was applied to remove noise before 8-bit transformation. The nuclei were segmented with the Intermodes thresholding method and smoothed with a median filter of radius = 4. For detection, the “Analyze Particles” algorithm was set to the following conditions: 0.20-4.00 µm^2^, circularity between 0.70 - 1.00, and including holes. The malaria pigment was segmented with the Moments thresholding method and detected with the “Analyze Particles” algorithm with the following conditions: 0.00-8.00 µm^2^, including holes. All the pigment ROI’s obtained were grouped into a single ROI, keeping track of the initial number of pigment spots. A smoothing/noise filter was omitted for the pigment because some spots are so small, or are too close together that the filter often renders them undetectable or merges them together.

To establish the above mentioned segmentation protocol, the conditions of the filters and the “Analyze Particles” algorithm were set with a random sample of 166 images of GFP-expressing or stained parasites (see Methods section). Up to 13 different segmentation methods were also tested to isolate the parasites, the nuclei, and the pigment form the rest of the image (Supplementary Figs.S3). The selection of a segmentation method was pondered by the following criterion: (1) major number of successful segmentations, (2) less bias in the segmentation between differentiation stages, and (3) less bias in the measurements of the segmented region of interest (ROI) relative to the manual segmentation.

Several attempts were made to segment the parasites from the bright field images; however successful segmentations were below 50%. This was mainly because of the Extracelular Matrix gel coating needed for purification and oocyst culture, which in occasions was not smooth enough introducing artifacts in the images. Therefore, the parasite segmentation was performed on the fluorescence images.

**Supplementary Figure 2:**
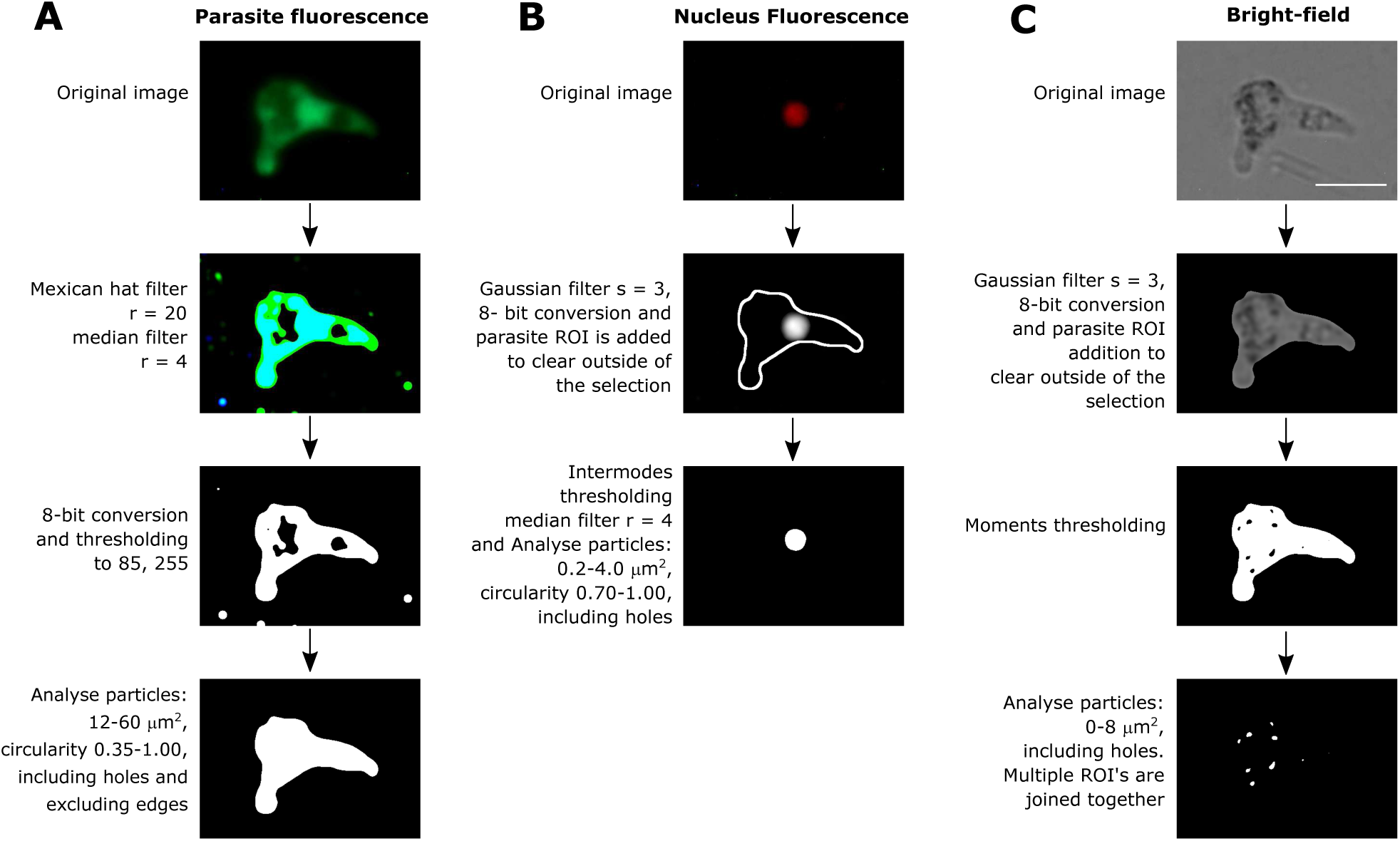
Procedures for images segmentation. Summary of the filters and operations applied to the images for segmenting the parasite **(A)**, the nucleus **(B)**, and pigment granules **(C)**. Only some representative images of the segmentation steps are shown.

**Supplementary Figure 3:**
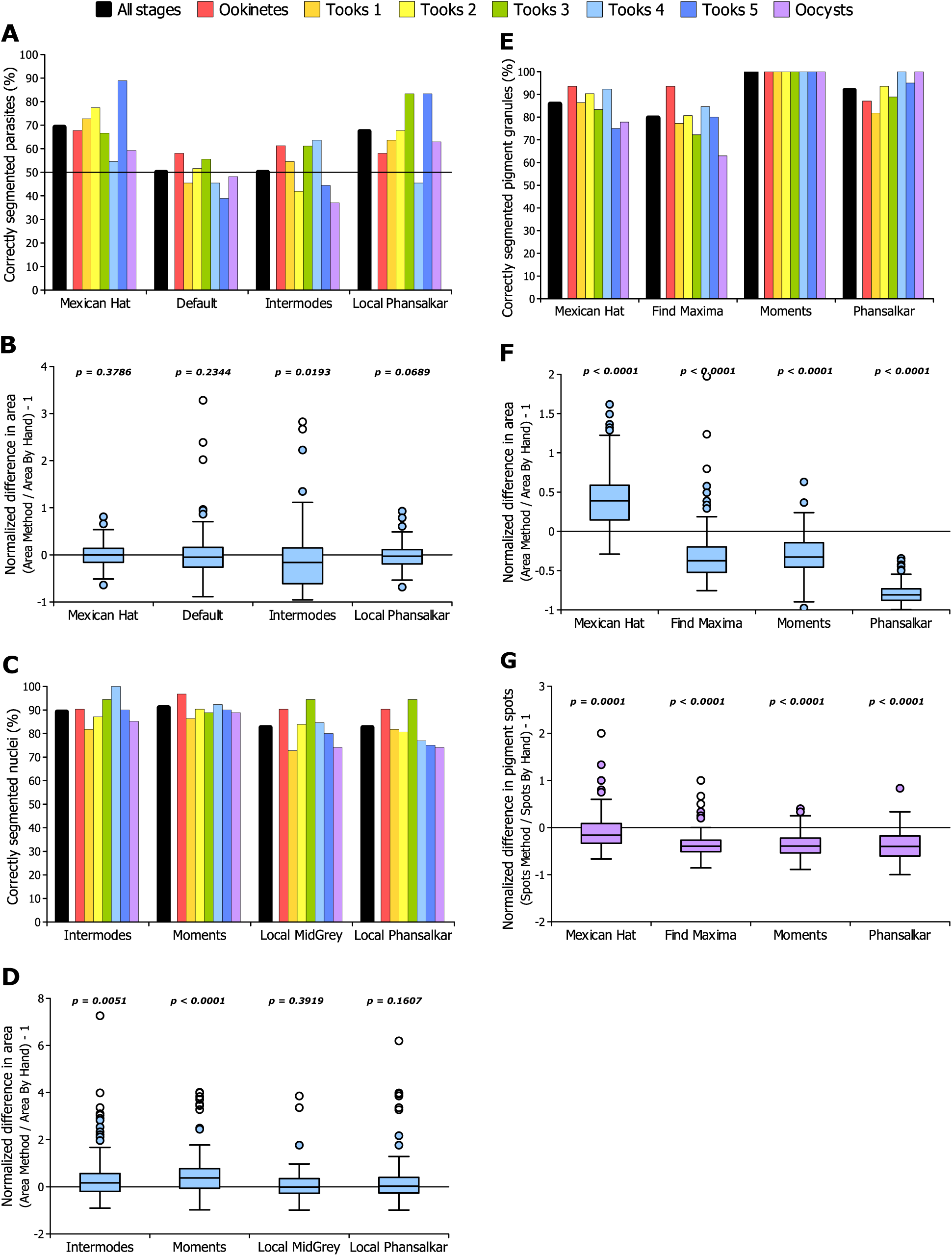
Four of the best methods tested for segmenting the parasites, the nuclei and the pigment granules. Out of 13 segmentation methods tested, here are represented four that performed the best in terms of the percentage of successfull segmentations, and the accuracy of the measurements compared to segmentations performed by hand. *(A)* To segment the parasite, the Mexican Hat filter was chosen for its implementation in the macro. *(B)* The Mexican Hat filter also was the one that produced the most accurate segmentation in regards to the area. *(C)* Despite the Moments thresholding was able to segment a slightly higher percentage of nuclei, it was also the one that differed the most in comparison to the segmentation done by hand in terms of the area of the nuclei *(D)*. Therefore, the Intermodes thresholding was instead chosen, which segmented about 90% of the images with good accuracy. *(E)* The Moments thresholding was the one chosen since it was able to segment 100% of the images despite not having a great accuracy *(F)* and *(G)*. Actually, almost all methods produced measurements that are substantially off of what was obtained by hand. *(B)* Data analyzed with Student’s *t* -test, comparing a given segmentation method vs. the segmentation done by hand. The horizontal line in (A) represents the minimum tolerable success where only 50% of the parasites were successfully segmented. *(D),(F)*, and *(G)* Data analyzed with Wilcoxon-Mann-Whitney’s test comparing a given segmentation method vs. the segmentation done by hand.

### Image processing for measurement of descriptors

The morphological characteristics depicted in Fig. 1 were regarded as measurable descriptors of the differentiation process. A total of 25 measurements were found to cover broadly these characteristics. Starting from the segmented parasite, nucleus, and pigment, the images were further processed to obtain the descriptors as follows. As seen in the Supplementary Fig. S4, the area, perimeter, adjustment to a circle, roundness, solidity (convex hull), aspect ratio (AR), adjustment to an ellipse, Feret’s diameter, and fitted bounding box were directly measured from the segmented image of the parasite using the ImageJ’s measure command. For the nuclei and pigment granules, the area was also measured from their corresponding segmented images.

The skeleton image was obtained from the segmented image of the parasite using the default “Skeletonize” algorithm built-in in ImageJ. The skeleton was detected with the “Analyze Particles” algorithm set without any restriction and measured. Since the skeleton is a one wide pixel line, the length of the skeleton was calculated by dividing the perimeter by two. Then the “Binary Connectivity” algorithm from the Morphological Operators package [2] was applied to the skeleton to obtain the number of branches and the number of junctions. This algorithm replaces the value of the pixels in a binary image depending on the number of foreground neighbors a pixel has. The resulting image after running the algorithm was thresholded to 1, 2 to only preserve the number of branches, which were counted with the “Analyze Particles” algorithm set with any restriction. For counting the number of junctions, the procedure was the same but thresholding the resulting image to 4, 5. These images were also used to determine the distance from the parasite’s center to the average tips of the branches and to the junction.

The relative position of the nucleus and pigment granules was measured by overlapping the ROI’s of these structures on an Euclidean distance map obtained from the segmented image of the parasite, where the foreground pixels are replaced with a gray value equal to the nearest background pixel. Measuring the average pixel intensity within the boundaries of the overlapped ROI gives the relative position, where greater values mean further from the center of the parasite. The number of pigment spots was calculated simply by counting the number of ROI’s generated after the “Analyze Particles” algorithm was run on the segmented image of the pigment.

**Supplementary Figure 4:**
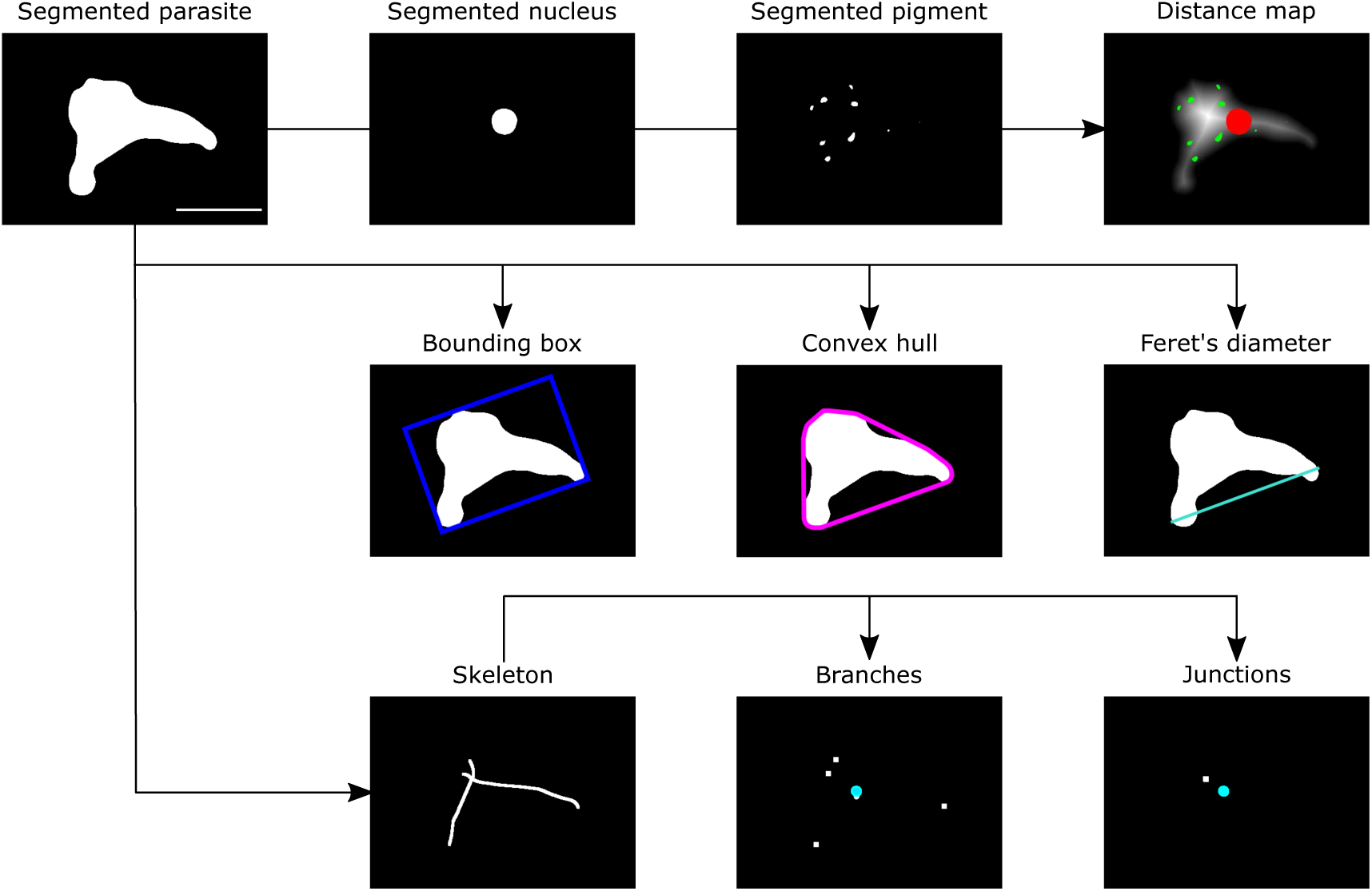
Image processing for the measurement of descriptors. From the segmented images, the area, perimeter, circularity, roundness, adjustment to an ellipse, and number of pigment spots are measured. The segmented parasite image also served to obtain the fitted bounding box, the convex hull and the Feret’s diameter. Then the skeleton of the parasite is obtained and further processed to count the number of branches and junctions (The images shown as example of the skeleton and the branches and junction were dilated for ease of visualization). The white squares represent the branches and the junction, while the round dot represent the average position of the branches. The cyan dot in the branches and junctions images represent the parasite center. For measuring the nucleus and pigment granules position, a distance map of the parasite was obtained and the nucleus (green) and pigment granules (red) ROI’s overlapped. For measuring the relative position of the nucleus, the distance map image was inverted in order to obtain increasing values as a representation of how far these structures are from the center of the parasite.

Supplementary Figures S5 to S7 contain the descriptive statistics for all the descriptors represented as violin plots. Table S1 contains the inferential statistics (*p* values) that denote the differences between the stages.

**Supplementary Figure 5:**
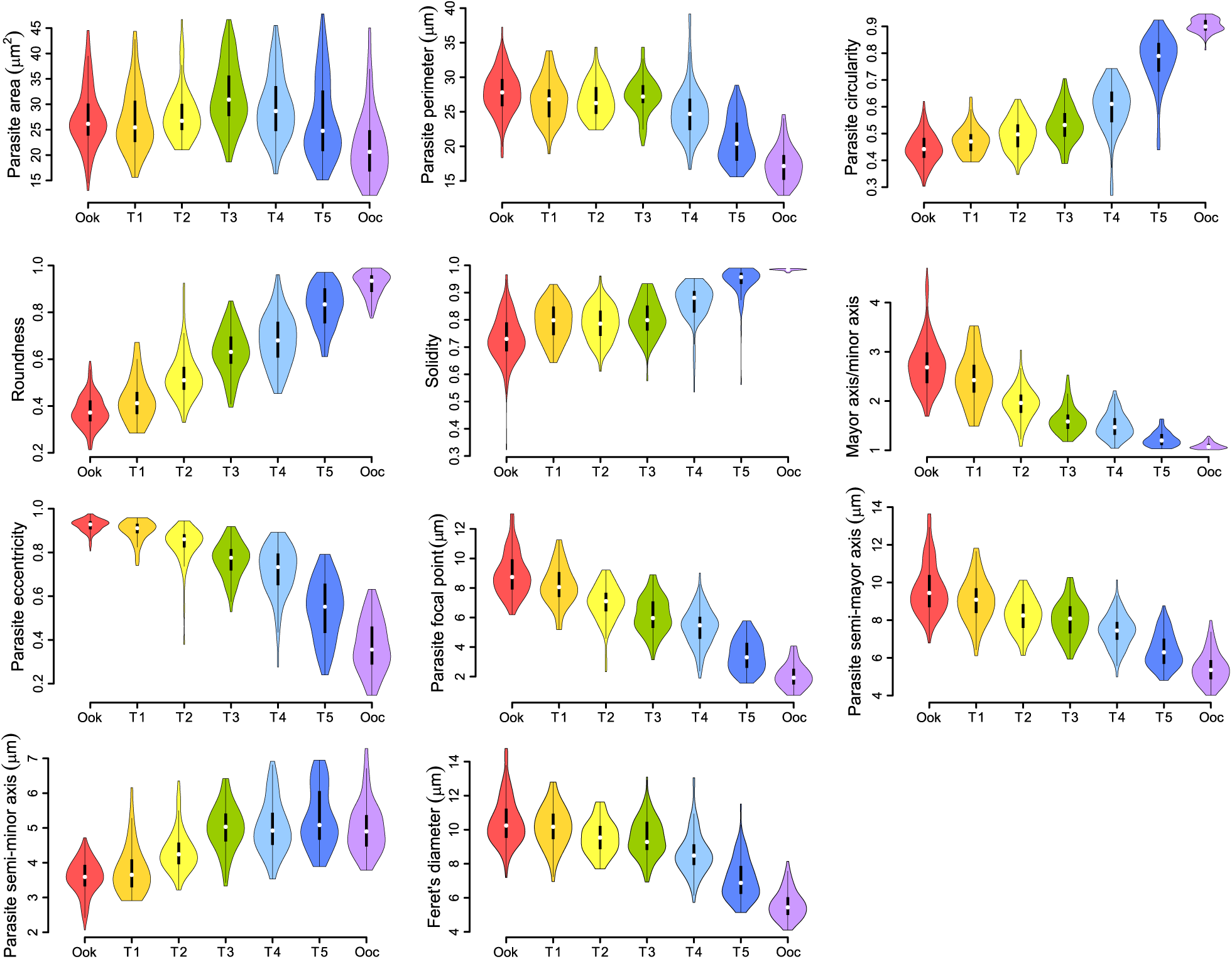
Descriptive statistics of the morphometric measurements of the differentiation stages. Violin plots representing the data of 706 automatically measured parasites (218 ookinetes, 68 tooks 1, 78 tooks 2, 71 tooks 3, 94 tooks 4, 92 tooks 5, and 85 oocysts). The white dots represent the median, the black bar the interquartile range, the black lines the upper and lower values, and the width of the curved lines the frequency distribution.

**Supplementary Figure 6:**
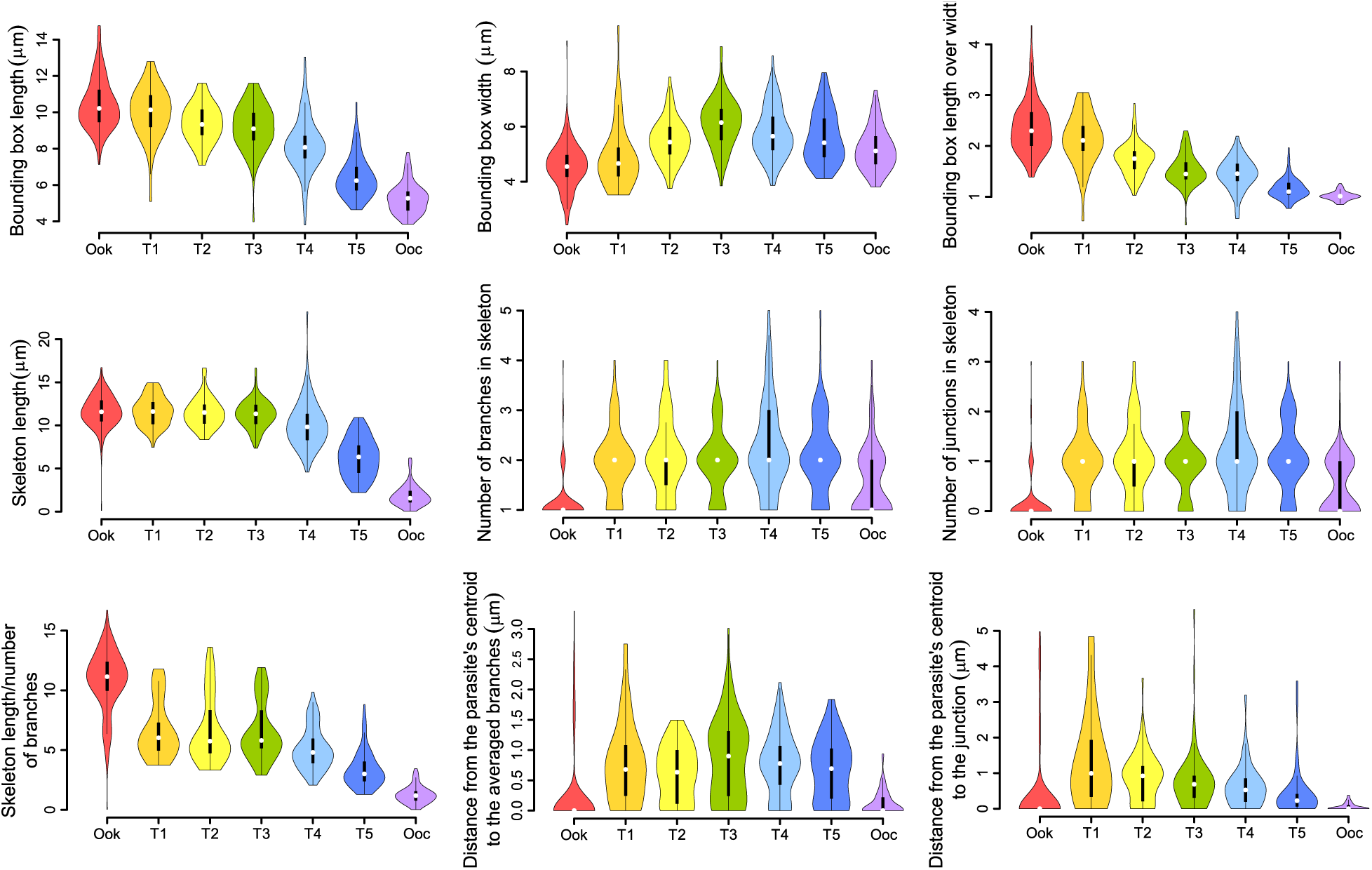
Descriptive statistics of the morphometric measurements of the differentiation stages. Violin plots representing the data of 706 automatically measured parasites (218 ookinetes, 68 tooks 1, 78 tooks 2, 71 tooks 3, 94 tooks 4, 92 tooks 5, and 85 oocysts). The white dots represent the median, the black bar the interquartile range, the black lines the upper and lower values, and the width of the curved lines the frequency distribution.

**Supplementary Figure 7:**
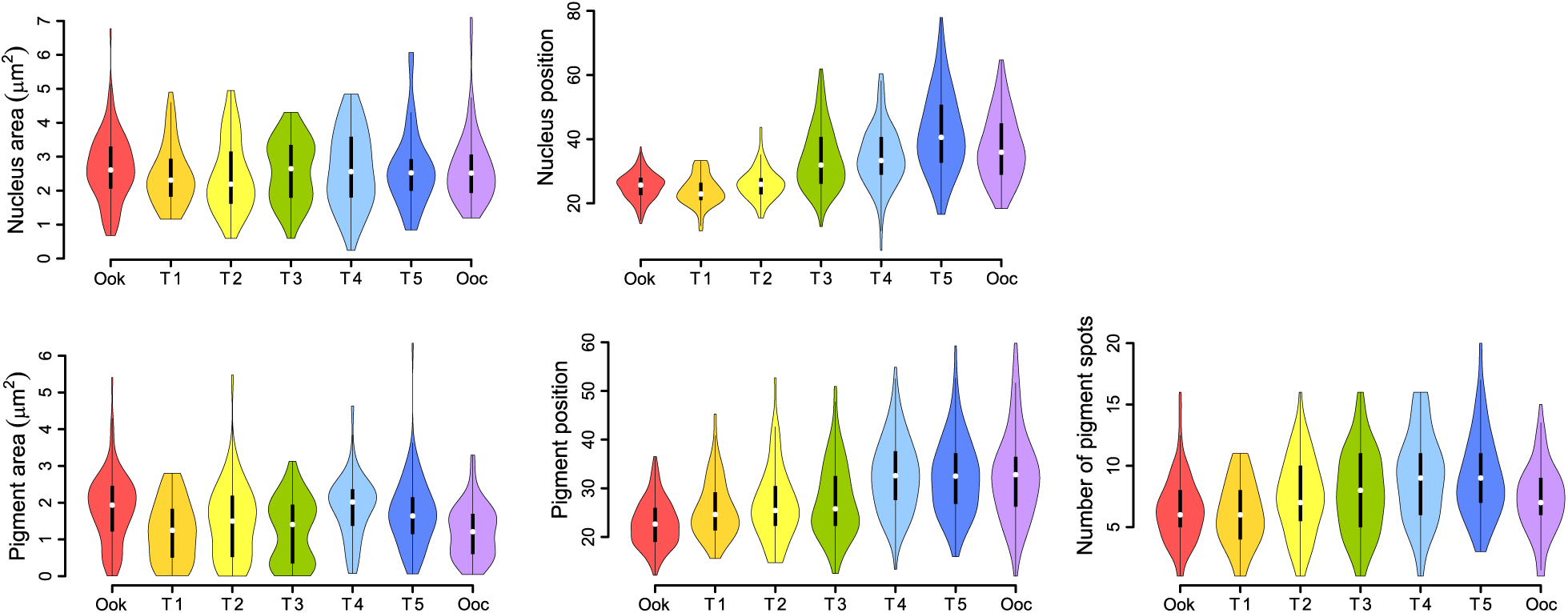
Descriptive statistics of the morphometric measurements of the differentiation stages. Violin plots representing the data of 706 automatically measured parasites (218 ookinetes, 68 tooks 1, 78 tooks 2, 71 tooks 3, 94 tooks 4, 92 tooks 5, and 85 oocysts). The white dots represent the median, the black bar the interquartile range, the black lines the upper and lower values, and the width of the curved lines the frequency distribution.

**Supplementary Table S1.** p-values of descriptor comparisons between differentiation stages analyzed by Kruskal-Wallis followed by Dunn’s test with Holm’s correction.

| Stage comparison | Parasite |  |  |  |  |  |  |  |  |  |  |  |  |  |
| --- | --- | --- | --- | --- | --- | --- | --- | --- | --- | --- | --- | --- | --- | --- |
|  | Area | Perimeter | Circularity | Roundness | Solidity | AR | Eccentricity | Focal Point | Semi-major axis | Semi-minor axis | Feret's diameter | Bounding box |  |  |
|  |  |  |  |  |  |  |  |  |  |  |  | Length | Width | Length/width |
| Ook-T1 | 1.0000 | 0.3141 | 0.1161 | 0.1124 | 0.0052 | 0.1134 | 0.1135 | 0.0599 | 0.1389 | 1.0000 | 0.8421 | 0.5935 | 0.5504 | 0.2047 |
| Ook-T2 | 0.9316 | 0.1874 | 0.0007 | <0.0001 | 0.0053 | <0.0001 | <0.0001 | <0.0001 | <0.0001 | <0.0001 | 0.0017 | 0.0012 | <0.0001 | <0.0001 |
| Ook-T3 | <0.0001 | 0.8852 | <0.0001 | <0.0001 | <0.0001 | <0.0001 | <0.0001 | <0.0001 | <0.0001 | <0.0001 | 0.0003 | <0.0001 | <0.0001 | <0.0001 |
| Ook-T4 | 0.1250 | <0.0001 | <0.0001 | <0.0001 | <0.0001 | <0.0001 | <0.0001 | <0.0001 | <0.0001 | <0.0001 | <0.0001 | <0.0001 | <0.0001 | <0.0001 |
| Ook-T5 | 0.8651 | <0.0001 | <0.0001 | <0.0001 | <0.0001 | <0.0001 | <0.0001 | <0.0001 | <0.0001 | <0.0001 | <0.0001 | <0.0001 | <0.0001 | <0.0001 |
| Ook-Ooc | <0.0001 | <0.0001 | <0.0001 | <0.0001 | <0.0001 | <0.0001 | <0.0001 | <0.0001 | <0.0001 | <0.0001 | <0.0001 | <0.0001 | <0.0001 | <0.0001 |
| T1-T2 | 1.0000 | 0.9637 | 0.1952 | 0.0055 | 0.8268 | 0.0054 | 0.0054 | 0.0243 | 0.0360 | 0.0024 | 0.1303 | 0.1665 | 0.0004 | 0.0056 |
| T1-T3 | 0.0005 | 0.8407 | 0.0066 | <0.0001 | 0.9692 | <0.0001 | <0.0001 | <0.0001 | 0.0015 | <0.0001 | 0.1013 | 0.0576 | <0.0001 | <0.0001 |
| T1-T4 | 0.1913 | 0.0496 | <0.0001 | <0.0001 | 0.0006 | <0.0001 | <0.0001 | <0.0001 | <0.0001 | <0.0001 | <0.0001 | <0.0001 | <0.0001 | <0.0001 |
| T1-T5 | 0.6427 | <0.0001 | <0.0001 | <0.0001 | <0.0001 | <0.0001 | <0.0001 | <0.0001 | <0.0001 | <0.0001 | <0.0001 | <0.0001 | <0.0001 | <0.0001 |
| T1-Ooc | 0.0005 | <0.0001 | <0.0001 | <0.0001 | <0.0001 | <0.0001 | <0.0001 | <0.0001 | <0.0001 | <0.0001 | <0.0001 | <0.0001 | 0.1403 | <0.0001 |
| T2-T3 | 0.0249 | 0.8746 | 0.1136 | 0.0316 | 0.9335 | 0.0319 | 0.0317 | 0.0748 | 0.2954 | <0.0001 | 0.8970 | 0.5716 | 0.0071 | 0.0982 |
| T2-T4 | 1.0000 | 0.0305 | <0.0001 | 0.0005 | <0.0001 | 0.0005 | 0.0005 | 0.0003 | 0.0021 | <0.0001 | 0.0018 | 0.0007 | 0.4659 | 0.0053 |
| T2-T5 | 0.3256 | <0.0001 | <0.0001 | <0.0001 | <0.0001 | <0.0001 | <0.0001 | <0.0001 | <0.0001 | <0.0001 | <0.0001 | <0.0001 | 0.8028 | <0.0001 |
| T2-Ooc | <0.0001 | <0.0001 | <0.0001 | <0.0001 | <0.0001 | <0.0001 | <0.0001 | <0.0001 | <0.0001 | 0.0001 | <0.0001 | <0.0001 | 0.1910 | <0.0001 |
| T3-T4 | 0.2174 | <0.0001 | 0.0174 | 0.1733 | 0.0011 | 0.1736 | 0.1746 | 0.1017 | 0.0372 | 1.0000 | 0.0014 | 0.0020 | 0.1666 | 0.2908 |
| T3-T5 | <0.0001 | <0.0001 | <0.0001 | <0.0001 | <0.0001 | <0.0001 | <0.0001 | <0.0001 | <0.0001 | 1.0000 | <0.0001 | <0.0001 | 0.0073 | <0.0001 |
| T3-Ooc | <0.0001 | <0.0001 | <0.0001 | <0.0001 | <0.0001 | <0.0001 | <0.0001 | <0.0001 | <0.0001 | 1.0000 | <0.0001 | <0.0001 | <0.0001 | <0.0001 |
| T4-T5 | 0.0117 | <0.0001 | 0.0003 | 0.0011 | 0.0002 | 0.0011 | 0.0011 | 0.0004 | 0.0013 | 1.0000 | 0.0001 | <0.0001 | 0.4039 | 0.0003 |
| T4-Ooc | <0.0001 | <0.0001 | <0.0001 | <0.0001 | <0.0001 | <0.0001 | <0.0001 | <0.0001 | <0.0001 | 0.9936 | <0.0001 | <0.0001 | 0.0016 | <0.0001 |
| T5-Ooc | 0.0003 | 0.0161 | 0.0103 | 0.0552 | 0.0057 | 0.0552 | 0.0549 | 0.0634 | 0.0414 | 1.0000 | 0.0123 | 0.0727 | 0.0782 | 0.0927 |

**Supplementary Table S1 (*continued*).** p-values of descriptor comparisons between differentiation stages analyzed by Kruskal-Wallis followed by Dunn's test with Holm's correction.
| Stage<br>comparison | Parasite |  |  |  |  |  | Nucleus |  | Pigment |  |  |
| --- | --- | --- | --- | --- | --- | --- | --- | --- | --- | --- | --- |
|  | Skeleton |  |  |  | Distance from<br>parasite's centroid to: |  | Area | Position | Area | Position | Spots<br>Number |
|  | Length | Branches | Junctions | Length/branches | Branches | Junction |  |  |  |  |  |
| Ook-T1 | 0.8637 | <0.0001 | <0.0001 | <0.0001 | <0.0001 | <0.0001 | 1.0000 | 0.4694 | <0.0001 | 0.0621 | 1.0000 |
| Ook-T2 | 1.0000 | <0.0001 | <0.0001 | <0.0001 | <0.0001 | <0.0001 | 0.5855 | 0.5081 | 0.0162 | 0.0010 | 0.0623 |
| Ook-T3 | 1.0000 | <0.0001 | <0.0001 | <0.0001 | <0.0001 | <0.0001 | 1.0000 | <0.0001 | <0.0001 | <0.0001 | 0.0039 |
| Ook-T4 | <0.0001 | <0.0001 | <0.0001 | <0.0001 | <0.0001 | <0.0001 | 1.0000 | <0.0001 | 1.0000 | <0.0001 | <0.0001 |
| Ook-T5 | <0.0001 | <0.0001 | <0.0001 | <0.0001 | <0.0001 | <0.0001 | 1.0000 | <0.0001 | 0.7422 | <0.0001 | <0.0001 |
| Ook-Ooc | <0.0001 | 0.0001 | 0.0005 | <0.0001 | 1.0000 | 1.0000 | 1.0000 | <0.0001 | <0.0001 | <0.0001 | 0.1171 |
| T1-T2 | 1.0000 | 1.0000 | 0.9520 | 1.0000 | 1.0000 | 1.0000 | 1.0000 | 0.3640 | 1.0000 | 1.0000 | 0.1289 |
| T1-T3 | 1.0000 | 0.9585 | 1.0000 | 1.0000 | 1.0000 | 1.0000 | 1.0000 | <0.0001 | 1.0000 | 1.0000 | 0.0343 |
| T1-T4 | 0.0038 | 1.0000 | 1.0000 | 0.0392 | 1.0000 | 0.6720 | 1.0000 | <0.0001 | 0.0003 | <0.0001 | <0.0001 |
| T1-T5 | <0.0001 | 1.0000 | 1.0000 | <0.0001 | 0.9518 | 0.0062 | 1.0000 | <0.0001 | 0.0479 | <0.0001 | <0.0001 |
| T1-Ooc | <0.0001 | 0.0129 | 0.0023 | <0.0001 | <0.0001 | <0.0001 | 1.0000 | <0.0001 | 0.8825 | 0.0001 | 0.1667 |
| T2-T3 | 1.0000 | 1.0000 | 1.0000 | 0.9620 | 1.0000 | 1.0000 | 1.0000 | <0.0001 | 1.0000 | 1.0000 | 1.0000 |
| T2-T4 | 0.0091 | 1.0000 | 1.0000 | 0.0456 | 1.0000 | 1.0000 | 1.0000 | <0.0001 | 0.0256 | <0.0001 | 0.1274 |
| T2-T5 | <0.0001 | 1.0000 | 1.0000 | <0.0001 | 1.0000 | 0.0265 | 1.0000 | <0.0001 | 0.9119 | 0.0001 | 0.1254 |
| T2-Ooc | <0.0001 | 0.0027 | 0.0005 | <0.0001 | <0.0001 | <0.0001 | 1.0000 | <0.0001 | 0.7860 | 0.0006 | 1.0000 |
| T3-T4 | 0.0103 | 1.0000 | 1.0000 | 0.0355 | 1.0000 | 0.7212 | 1.0000 | 0.7779 | <0.0001 | 0.0001 | 0.2074 |
| T3-T5 | <0.0001 | 1.0000 | 1.0000 | <0.0001 | 1.0000 | 0.1323 | 1.0000 | 0.0016 | 0.0425 | 0.0003 | 0.1942 |
| T3-Ooc | <0.0001 | 0.0027 | 0.0005 | <0.0001 | <0.0001 | <0.0001 | 1.0000 | 0.4788 | 1.0000 | 0.0015 | 1.0000 |
| T4-T5 | <0.0001 | 1.0000 | 1.0000 | 0.0021 | 1.0000 | 0.2686 | 1.0000 | 0.0256 | 0.8492 | 0.8031 | 0.9784 |
| T4-Ooc | <0.0001 | <0.0001 | <0.0001 | <0.0001 | <0.0001 | <0.0001 | 1.0000 | 0.9312 | <0.0001 | 1.0000 | 0.0314 |
| T5-Ooc | 0.0093 | 0.0001 | <0.0001 | 0.0018 | <0.0001 | 0.0003 | 0.9679 | 0.4219 | 0.0091 | 1.0000 | 0.0322 |

Ook = ookinete, T1 = took 1, T2 = took 2, T3 = took 3, T4 = took 4, T5 = took 5, Ooc = oocyst

### Automatic classification of the oocyst differentiation stages

A set of 28 parasites (4 for each stage) with their descriptors computed as in Fig.2 were visually classified and used as seed to build a training database for the principal component analysis (PCA). The PCA successfully spread the data of the parasites in a multidimentional space, where 80% of the variation was explained by the first 4 PC’s and *>*95% of the variation was explained by the first 10 PC’s. For predicting the stage of a parasite, the first 10 PC’s were used. First, the centroids of the data clusters for each stage were calculated as the mean of the coordinates for that stage:

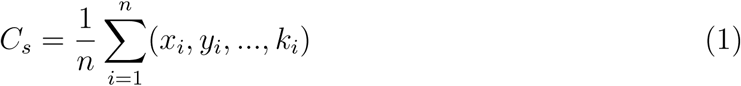

where *C*_*s*_ is the centroid of a stage, *n* is the number of parasites in that stage, and *x*_*i*_, *y*_*i*_, …, *k*_*i*_ are the 10 coordinates of each parasite. Then the PC’s of a parasite-to-classify were predicted, and the Euclidean distance to a centroid for each stage was calculated with the generalization of the Pythagorean Theorem equation:

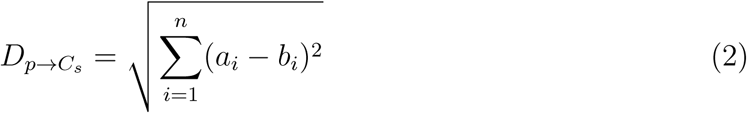

Where 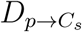 is the distance between the PC’s of the parasite-to-classify and the centroid of a stage, *a*_*i*_ are the coordinates of the parasite-to-classify, and *b*_*i*_ are the coordinates of the centroid of a stage, both for the *n* dimensions. Once the distance of the parasite-to-classify to each centroid was calculated, the algorithm returned the minimum distance to a centroid as the corresponding predicted stage of the parasite. That is, the closest centroid of a data cluster representing a stage to the parasite’s predicted PC’s, is assigned as its differentiation stage (Supplementary Fig. S8). Once a parasite has been classified, its measurements are included in the training database to extend it only if this option has been enabled.

**Supplementary Figure 8:**
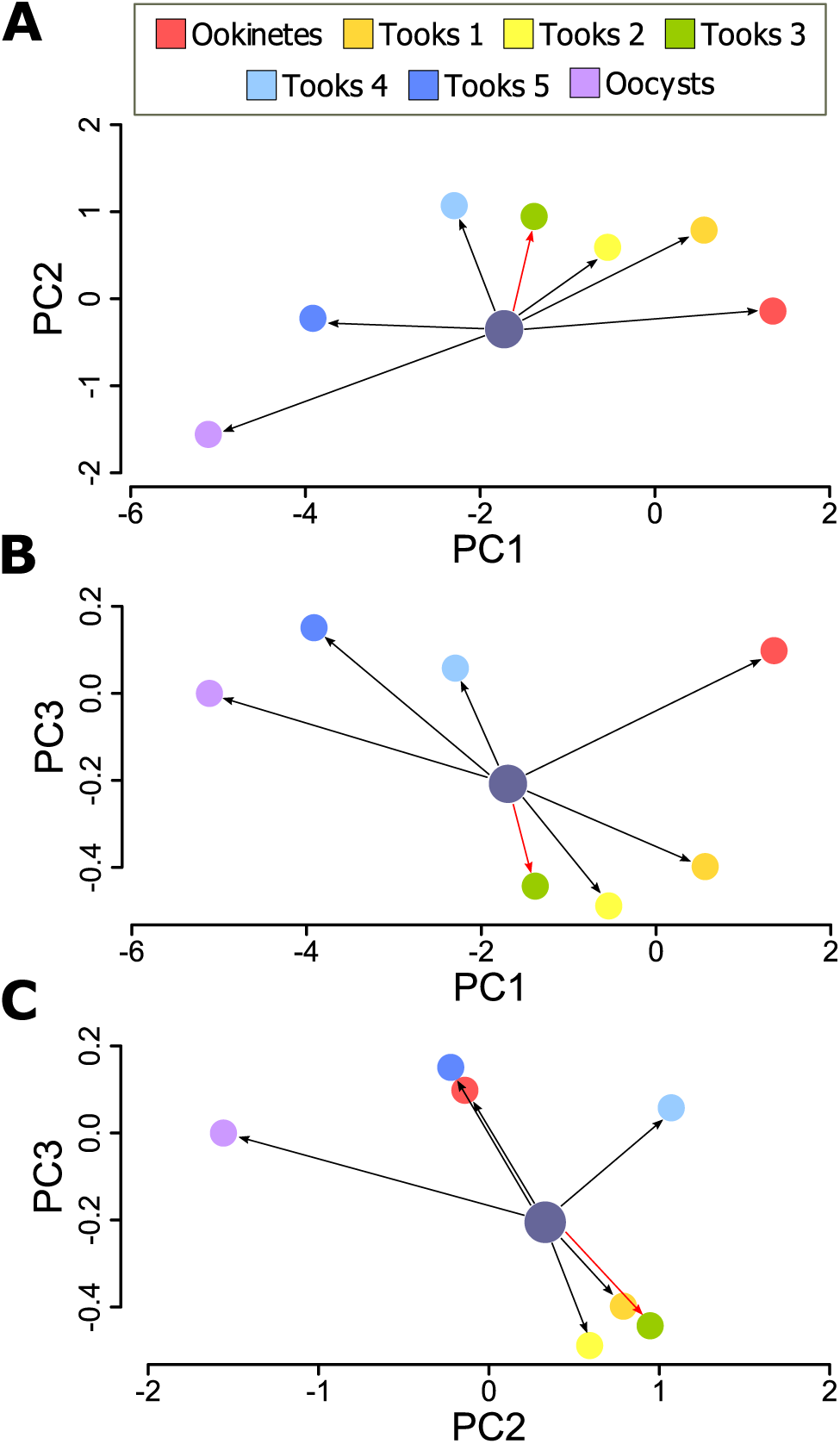
Centroids of the data clusters for each differentiation stage in the first three PC’s. **(A)** PC’s 1 and 2. **(B)** PC’s 1 and 3, and **(C)** PC’s 2 and 3. For classification, the distance from the parasite-to-classify (here represented as a big metallic blue circle) to each centroid (represented as colored smaller circles) considering the first 10 PC’s is calculated, returning the closest centroid as the assigned stage of the parasite-to-classify. In this example the parasite was classified as a took 3 (red arrow).

For evaluating the performance of the classification algorithm, the precision, recall and Jaccard indices were calculated during training (Supplementary Figure S9). These indices started form a value of 1 because of the 28 parasites used as seed. Then, these values started to decrease as the algorithm entered a phase where the centroids for each stage get defined in space. As more parasites entered the training database, the values bottomed to a minimum and slowly started to increase again, meaning that the algorithm, between the 400^*th*^ and 500^*th*^ parasite, got better at classifying as more parasites entered the database.

**Supplementary Figure 9:**
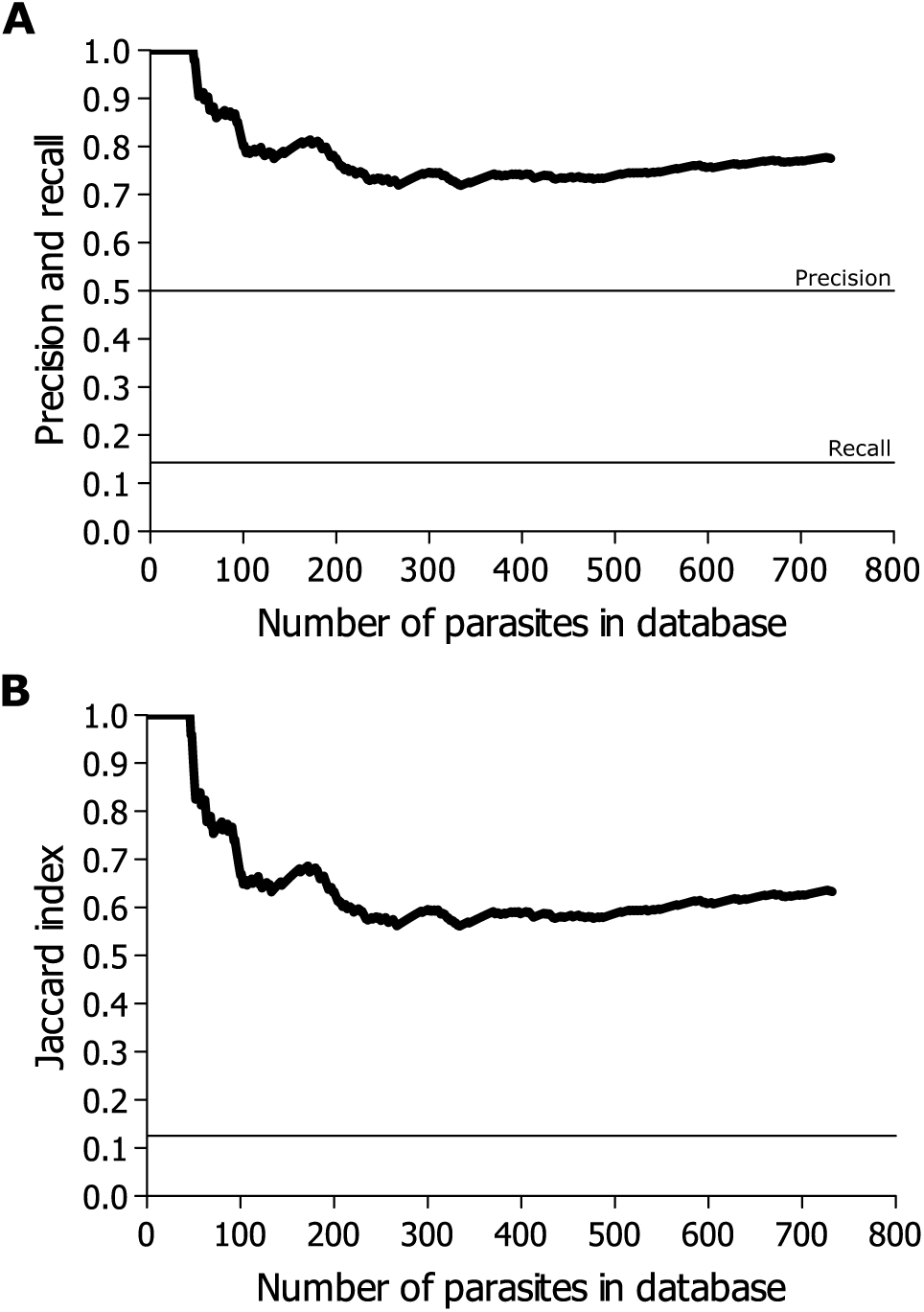
Performance evaluation indices for the classification. **(A)** Precision and recall of the classifications of all stages as the database size increased. Since the wrong assignments for a stage implies wrong assignments for another stage, the precision and recall behaved equally when all stages were considered. **(B)** Jaccard index of the classifications of all stages as the database size increased. The horizontal lines represent the value at which the classification would happen at random. Below the lines, the parasites would be systematically misassigned. The horizontal lines take different values because the probability of obtaining a true positive (TP), a false negative (FN), and a false positive (FP) are different. The probability of obtaining stage *x* at random when it is actually stage *x* (that is a TP) is 1*/*7, while the probability of obtaining any other stage (a FN) is 6*/*7. The probability of obtaining stage *x* when it is actually any other stage is also 1*/*7 (a FP).

